# RNF13 is a novel interactor of iduronate 2-sulfatase that modifies its glycosylation and maturation

**DOI:** 10.1101/2025.06.20.660705

**Authors:** Valérie C. Cabana, Antoine Y. Bouchard, Audrey M. Sénécal, Laurent Cappadocia, Marc P. Lussier

## Abstract

Mucopolysaccharidosis type II, also known as Hunter syndrome, is a rare and fatal disease caused by mutations in the *iduronate 2-sulfatase (IDS)* encoding gene. The enzymatically inactive variant proteins lead to pathological accumulation of glycosaminoglycans in the lysosomes, causing dysfunction in multiple organs. IDS is expressed as a precursor protein, and its processing and lysosomal targeting are crucial for proper enzymatic activity. However, IDS intracellular dynamic is poorly understood and a better understanding of its processing mechanisms would benefit the development of new therapeutic strategies. AlphaFold 3 predicted an interaction between IDS and the E3 ubiquitin ligase RNF13. Co-immunoprecipitation assays confirm this interaction and further show that RNF13 interacts preferentially with a predominantly underglycosylated immature form of IDS, resulting in altered IDS glycosylation and maturation. The results demonstrate that IDS glycosylation site Asn246 is important for lysosomal targeting, although its glycosylation is not altered by RNF13. This study also unravels that RNF13 forms a heterodimer with the E3 ubiquitin ligase RNF167 that modify both RNF13 and RNF167 lysosomal trafficking. In addition, the heterodimer interacts and alters IDS differently than RNF13 or RNF167 alone. RNF13 catalytic E3 ligase activity is required to generate an underglycosylated form, but not that of RNF167. This study exposes that the proteasome rapidly degrades IDS underglycosylated forms, and RNF13 exerts a protective effect. Overall, this study reveals a novel and dual role of RNF13 on IDS maturation and degradation, providing mechanistic insights into IDS trafficking.

## INTRODUCTION

Mucopolysaccharidosis II (MPS II), also known as Hunter syndrome, is a rare lysosomal storage disease caused by mutations in the *iduronate-2-sulfatase (IDS)* gene. IDS is an enzyme required for the lysosomal degradation of heparan sulfate and dermatan sulfate of glycosaminoglycans (GAGs) [1]. Enzyme deficiency leads to the progressive accumulation of GAGs, resulting in multisystemic dysfunction, with approximately two-thirds of patients exhibiting neurological impairment [2]. While enzyme replacement therapy is currently the standard of care, it shows minimal to no effect on bones, heart or neurological symptoms, supporting the need for new therapeutic strategies [3].

IDS protein is expressed as a precursor protein of 550 amino acids, with a signal peptide, and eight N-linked glycosylation sites (Fig. 1A) [4]. Protein synthesis occurs in the endoplasmic reticulum (ER) as a 75 kDa glycosylated precursor, while removal of the glycosylation, an important mechanism in the proteasomal degradation of glycoproteins in the ER [5], would lead to an approximately 60 kDa IDS precursor form (Fig. 1B-C) [6]. During its maturation in the Golgi, it is subjected to further glycosylation and phosphorylation to create mannose-6-phosphate (M6P) moieties recognized by M6P receptors (M6PRs) in the trans-Golgi network (TGN), leading to a 90 kDa form (Fig. 1D) [6]. Subsequent lysosomal proteolytic cleavage separates the 55 kDa mature N-terminal SD1 and the 18 kDa C-terminal SD2 (Fig. 1E) [6]. Both domains are stably associated to form an enzymatically competent complex [4]. MPS II-causing variants have been reported to impair proteolytic cleavage, precluding IDS maturation and enzymatic activity [7, 8]. A variant with an attenuated phenotype of the disease was shown to regain some enzymatic activity if translocation to the lysosomes was improved [7]. However, our knowledge on the trafficking pathways regulating IDS lysosomal targeting is still limited, and a better understanding would benefit the development of new therapeutic target.

**Figure 1:**
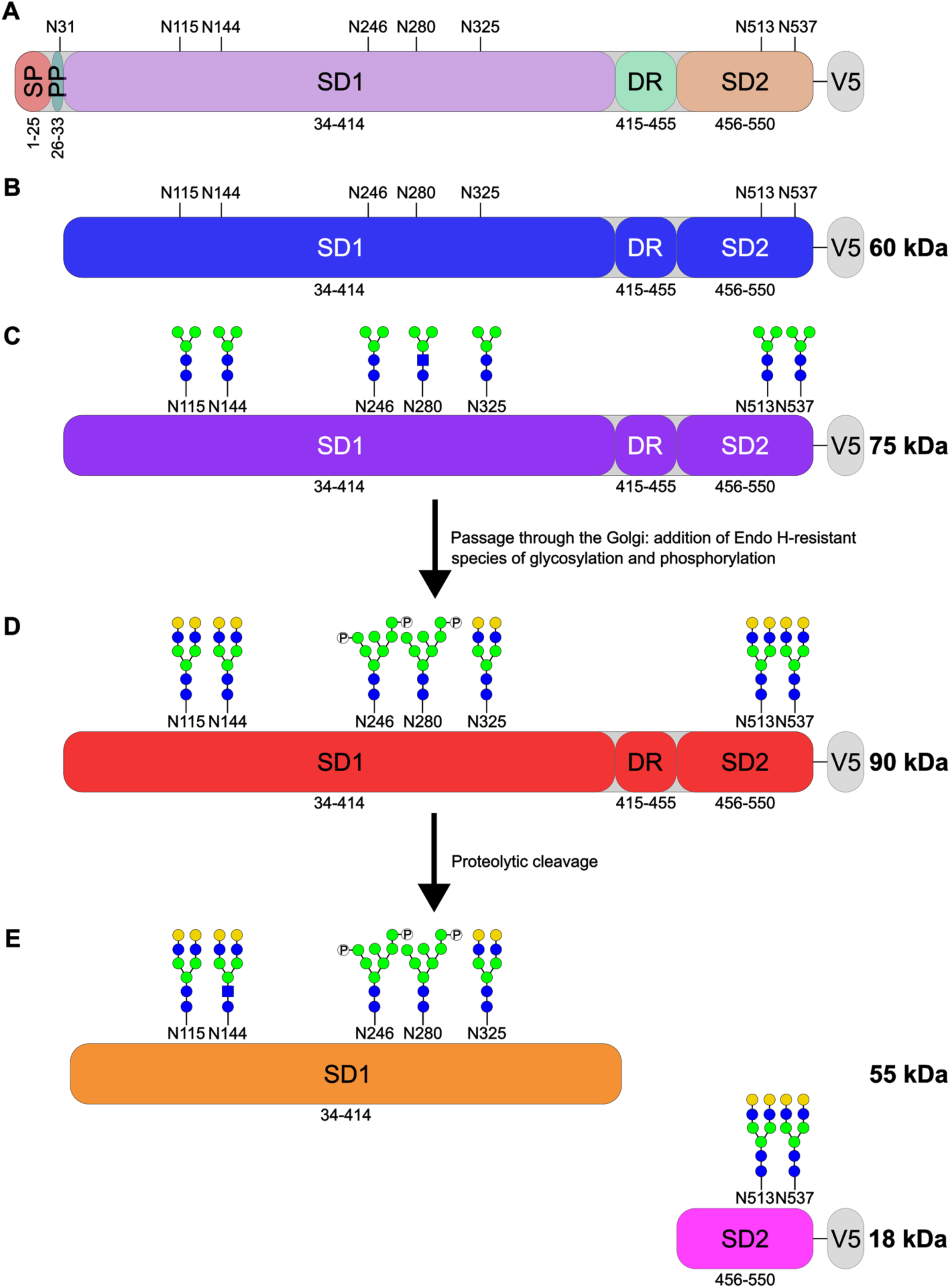
Schematic of IDS different forms based on the maturation and processing. (**A**) The complete protein is 550 amino acids long and contains a signal peptide (SP), a propeptide (PP), a SD1 domain, a disordered region (DR), a SD2 domain, and eight N-glycosylation sites. (**B-C**) The precursor form is (**B**) 60 kDa without any N-glycosylation (in blue) and (**C**) 75 kDa with the N-glycosylation added in the ER (in purple). (**D**) Further processing in the Golgi adds complex (N115, N144, N325, N513, N537) or high mannose types that are phosphorylated (N246, N280), leading to the 90 kDa form (in red). (**E**) The mature form is obtained after proteolytic cleavage, where the SD1 (in orange) and SD2 (in pink) domains are separated. (**A-E**) In this study, a V5 tag was added at the C-terminus and used to distinguish the different processed forms.

For soluble proteins such as IDS to be transported in the endolysosomal system, they need to be recognized by a specific receptor and bind to its luminal side. For instance, the luminal side of M6PRs recognize IDS glycosylation state, while the receptors bind and are trafficked by AP-1, one of the adaptor protein (AP) complexes, the main regulators of post-Golgi trafficking pathways [9]. Although it is not known if AP-1 regulates IDS trafficking, AP-1 and AP-3 are responsible for recognizing specific sorting signals on cytoplasmic tails of transmembrane proteins for late endosomal and lysosomal targeting [10, 11]. In addition, one of the subunit of the AP-3 complex is reported on the BioGRID database to interact with IDS [12, 13], but not AP-1, which could suggest an alternate and unknown trafficking pathway. Intriguingly, we have previously shown that trafficking of RNF13, an ubiquitin E3 ligase whose function is associated with ER stress, various form of autophagy, and protein degradation and localization [14-17], is regulated by AP-1 and AP-3 that recognizes motifs in its C-terminal [18, 19]. Furthermore, RNF13 variants that alter the lysosomal localization of the protein cause a severe neurodegenerative disease, with some symptoms overlapping with those of MPS II [18, 20, 21]. RNF13 possesses a luminal protease-associated (PA) domain, a transmembrane (TM) domain, and a cytosolic Really Interesting New Gene (RING) domain, making it a member of the PA-TM-RING family [22]. While multiple functions have been recently reported for its cytosolic E3 ligase function [15-17, 23], knowledge on the function of its luminal PA domain remains limited, except that the plant ortholog of RNF13 use this domain to bind soluble proteins in the ER and transport them to the vacuole [24].

In this study, AlphaFold 3 was used to predict whether RNF13 could be involved in IDS trafficking to the endolysosomal pathway. This study demonstrates, using co-immunoprecipitation assays, that RNF13 interacts with a precursor form of IDS that is predominantly deglycosylated. Unexpectedly, it reveals that RNF13 alters IDS glycosylation, changes its maturation, and increases the abundance of an underglycosylated precursor form. To control that IDS alterations caused by RNF13 are specific, a second PA-TM-RING E3 ligase, RNF167, was used. Both RNF13 and RNF167 exerts different effects on IDS, and the combination further increases the underglycosylated form of IDS. Our study reveals that RNF13 and RNF167 heterodimerize, but only the E3 ligase activity of RNF13 is required to induce the underglycosylated form, which is normally rapidly degraded by the proteasome but is protected by RNF13. Our study provides new insights into IDS maturation and opens up new avenues for a deeper understanding of MPS II pathophysiology.

## RESULTS

### RNF13 interacts with IDS, altering its glycosylation and maturation

To better understand IDS’s trafficking, we parsed the BioGRID database of protein-protein interaction and found that one of the AP-3 complex subunit is reported to interact with IDS [12, 13]. AP-3 is responsible for the trafficking of transmembrane proteins towards the lysosomes [25]. However, IDS is soluble in the ER lumen and would thus need to interact with a receptor. As the RNF13 plant ortholog is a reported sorting receptor that transports soluble proteins from the Golgi to the vacuole [24], we wondered if human RNF13 might mediate similar functions on IDS. Using AlphaFold 3, a predicted interaction was identified between the precursor forms of IDS and RNF13. The predicted model shows that RNF13 PA domain engages residues that are restricted to the SD1 portion of IDS (Fig. 2A). The aliphatic portion of IDS residues Pro243, Glu245 and Asn246 stack against the walls of a hydrophobic cavity formed, in part, by the aliphatic portions of Asn103, Phe104, and Ile130 of RNF13 (Fig. 2A). IDS Glu245 forms a hydrogen bond with the main chain amine of Arg99 while simultaneously forming salt bridges with the side chain of Arg99. IDS Asn246 interacts with the main chain of RNF13 Ser131 and Gly133. RNF13 Arg54 interacts with the main chain of Lys57 and Leu58 as well as the side chain of Arg60 in IDS. In turn, IDS Arg60 interacts with the side chain of RNF13 Ser131. Finally, RNF13 Asn135 forms hydrogen bonds with the main chain of Thr248 as well as the side chains of Thr248 and Glu245 in IDS (Fig. 2A). To better understand the consequences of IDS processing on this interaction, AlphaFold 3 was used with RNF13 and the N-terminal SD1 and C-terminal SD2 as separate proteins to mimic IDS enzymatically competent complex. This alteration changes the prediction to a model where RNF13 interacts with the hydrophobic portion of the SD2 fragment (Fig. 2B). Specifically, Ile36, Leu66 and the aliphatic parts of Tyr41 and Tyr43 of SD2 are embedded in a hydrophobic pocket of RNF13, mainly composed of the aliphatic parts of Phe41, Glu42, Tyr164, Glu165 and His169 (Fig. 2B). Additionally, the hydroxyl group of Tyr7 and Tyr41 of SD2 are both stabilized by Asn43 and Tyr164 of RNF13, respectively, the latter forming a hydrogen bond (Fig. 2B).

**Figure 2:**
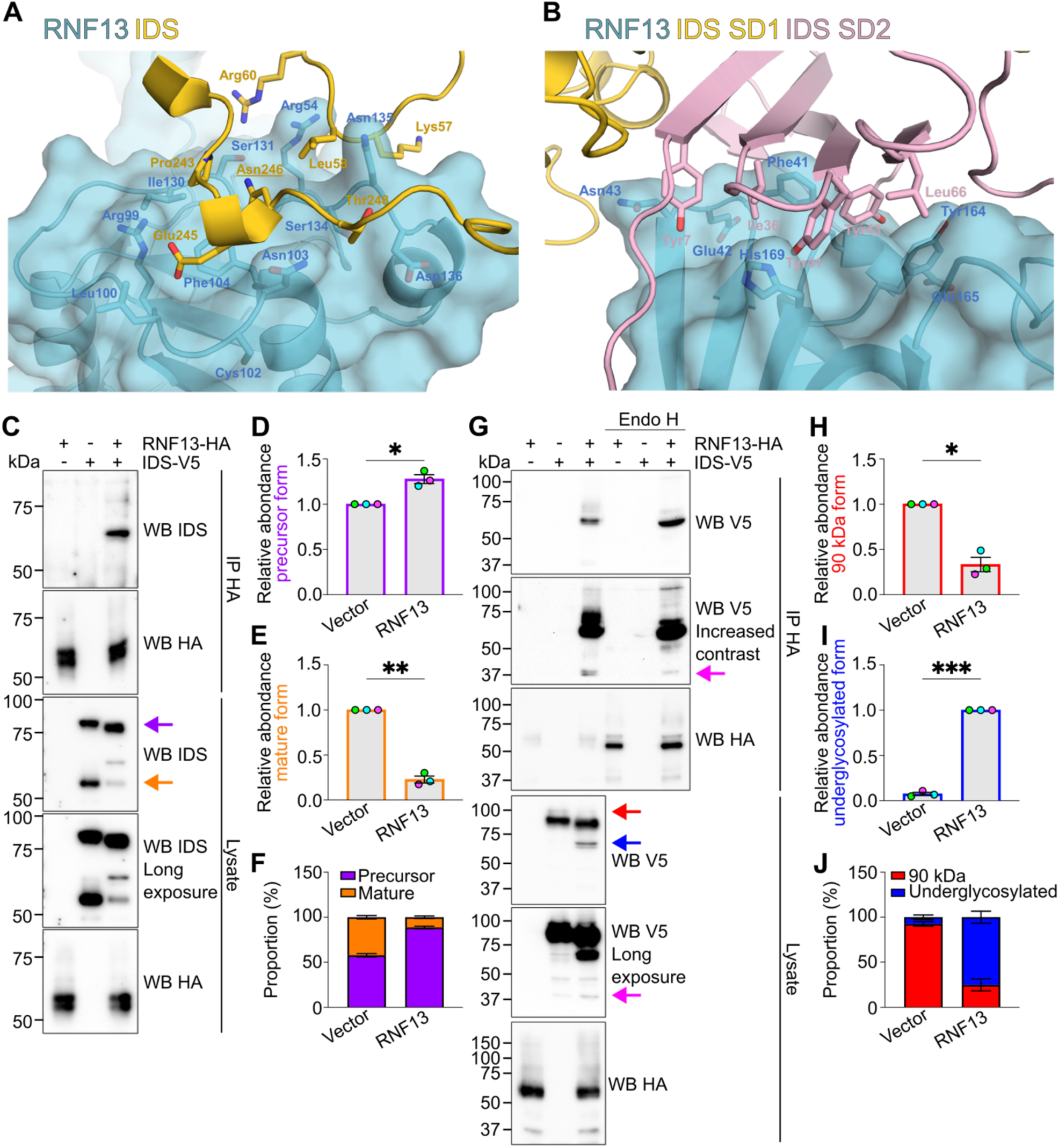
RNF13 interacts with precursor underglycosylated IDS and alters its processing. (**A-B**) AlphaFold 3 predicted an interaction between RNF13 (cyan) and (**A**) the precursor form of IDS (yellow) or (**B**) the SD2 portion of the cleaved form of IDS (pink). (**C, G**) Representative (N=3) immunoblots show IDS-V5 and RNF13-HA detection in HA-immunoprecipitated complexes from transfected HEK293T/17 cell lysates (**G**) digested with Endo H. The arrows represent the precursor (purple), mature (orange), 90 kDa (red), and underglycosylated (blue) forms, as well as the SD2 fragment (pink). (**D-E, H-I**) The strip plot represents the relative protein levels of IDS (**D**) precursor, (**E**) mature, (**H**) 90 kDa, and (**I**) underglycosylated form. (**F, J**) The summary bar plots represent the proportion of (**F**) precursor/mature and (**J**) 90kDa/underglycosylated form. All results are mean ± SEM from N=3. * *p* < 0.05; ** *p* < 0.01; *** *p* < 0.001 (two-tailed paired t test).

IDS/RNF13 interaction was experimentally validated using co-immunoprecipitation from HEK293T/17 cell lysates expressing RNF13-HA and IDS-V5 proteins. An anti-IDS immunoblot revealed the presence of a ∼65 kDa band in the anti-HA immunopurified protein complex (Fig. 2C). RNF13 exogenous expression led to important modifications in the IDS bands pattern in the lysate. Specifically, the abundance of IDS precursor glycosylated form is significantly increased while also presenting a slight change in molecular weight (MW) that could indicate the loss of a glycosylation site (purple arrow; Fig. 2C-D). In comparison, the mature form is significantly less abundant, while the MW is restablish to that of IDS alone (orange arrow; Fig. 2C, E). When comparing both forms, exogenous RNF13 increased IDS precursor proportion from 58% to 89% (Fig. 2F). The deglycosylated precursor form of IDS is around 60 kDa and some species of IDS glycosylated mature form have similar MW (smear extending up to ∼65 kDa; Fig. 2C) [6, 7]. To distinguish the different forms, a V5-tag was added onto the C-terminal, where only the mature forms will not be detected using an anti-V5 (Fig. 1E). To confirm which IDS form interacts with RNF13, co-immunoprecipitation experiments using anti-HA were performed from HEK293T/17 cell lysates (Fig. 2G). Endoglycosidase H (Endo H) digestion of the immunopurified complex revealed an IDS band, detected by anti-V5, around 62 kDa and a faint band around 68 kDa, while the non-digested sample showed two bands around 65 kDa and a smear extending up to 75 kDa, suggesting that it is predominantly deglycosylated but not completely (hereafter referenced as an underglycosylated form) precursor form containing the V5-tag. Unlike the subtle changes observed for IDS, RNF13 exhibited a sharp change in MW and band intensity (Fig. 2G), confirming efficient Endo H digestion as previously shown [19]. Interestingly, a small quantity of a C-terminal fragment of IDS is also detected in the immunopurified complex, even if this form is barely present in the lysate (pink arrows; Fig. 2G). Furthermore, IDS underglycosylated precursor form was barely detectable when IDS was expressed alone, in contrast to when exogenous RNF13 was present (Fig. 2C, G). While the underglycosylated form (blue arrow; Fig. 2G) increased significantly, the 90kDa form (red arrow; Fig. 2G) is less abundant upon RNF13 co-expression (Fig. 2H-J). Overall, our results suggest that RNF13 interacts with a predominantly underglycosylated precursor form of IDS and, to some extent, a C-terminus fragment, and alters IDS glycosylation and maturation.

### RNF13 trafficking to late endosomes/lysosomes is delayed by presence of IDS

It is established that IDS glycosylation is important for its lysosomal targeting because M6PR recognizes it [4, 26]. As RNF13 alters IDS glycosylation and reduces the abundance of the mature form, an immunofluorescence assay was performed on HEK293T/17 cells expressing RNF13-HA, IDS-V5 and GFP-Rab7 to evaluate if its lysosomal targeting was affected. First, ectopic expression of the late endosomal marker GFP-Rab7 did not alter the co-localization between IDS and RNF13 when compared to GFP alone (Fig. 3A-C). The images show that IDS is mostly found in the ER but also makes some contact with Rab7 (Fig. 3A). However, whereas RNF13 expression did not alter the Manders’ overlap coefficient (MOC) of IDS over Rab7 (Fig. 3D), IDS expression diminished the MOC of RNF13 over Rab7 (Fig. 3E). As the anti-IDS recognizes both the precursor and mature forms, the ER signal might be predominantly from the precursor form. To increase the detection of the lysosomal mature form, transfected HEK293T/17 cells were incubated with cycloheximide (CHX), a protein synthesis inhibitor, allowing the already synthesized proteins to mature [7] (Fig. 3F). The treatment decreased the overall fluorescence intensity without affecting the MOC between IDS and RNF13 (Fig. 3F-H). While a slight increase in MOC was observed for IDS over Rab7 following CHX treatment, RNF13 overexpression had no effect (Fig. 3I). However, when IDS was expressed alone, it exhibited a more punctate staining pattern after CHX treatment, a phenotype that was not observed with RNF13 (Fig. 3F). Notably, RNF13 alone colocalized with Rab7, while IDS overexpression decreased the MOC (Fig. 3E). CHX treatment rescued the colocalization between RNF13 and Rab7 (Fig. 3J), suggesting that IDS might delay RNF13 trafficking to late endosomes/lysosomes, which would be consistent with an interaction in the ER. Although it did not support our initial hypothesis where RNF13 could have been a lysosomal transporter for IDS, these results suggest that RNF13 interacts with a predominantly underglycosylated precursor form of IDS, thereby affecting its glycosylation, maturation, and trafficking.

**Figure 3:**
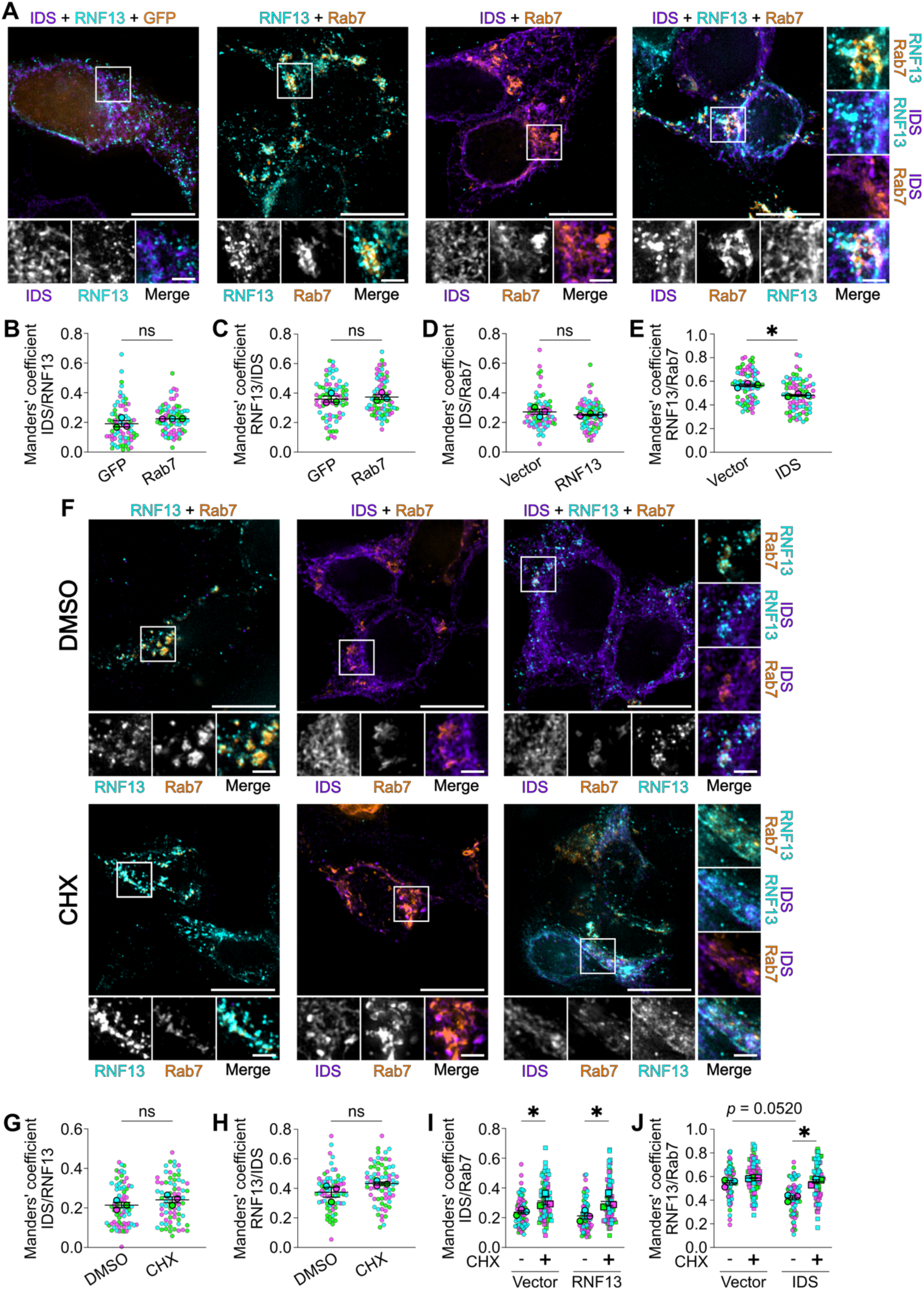
RNF13 trafficking to late endosomes/lysosomes is delayed by the presence of IDS. (**A, F**) Representative immunofluorescence images show HEK293T/17 cells expressing IDS-V5 (purple), RNF13-HA (cyan) and GFP-Rab7 (orange) (**F**) treated with CHX or vehicle for 4h30. Scale bars indicate 10 μm for whole cell image and 2 μm for higher magnification. (**B-E, G-J**) The jittered individual value plot represents MOC of (**B, G**) IDS over RNF13, (**C, H**) RNF13 over IDS and (**D, I**) IDS or (**E, J**) RNF13 over Rab7. All results are mean ± SEM from N=3. Each color represents an independent experiment (N=3). Smaller points represent individual cells analyzed (n=72). Bigger points represent the mean of an independent experiment. * *p* < 0.05; ** *p* < 0.01; *** *p* < 0.001 ((**B-E, G-H**) two-tailed paired t test or (**I-J**) two-way RM-ANOVA with multiple comparisons).

### IDS glycosylation site Asn246 is important for its maturation

The Asn246 amino acid of IDS is modified with M6P-containing glycans (Fig. 1D-E) [27]. Given the predicted binding of Asn246 with RNF13 (Fig. 2A), it is hypothesized that the interaction between the two proteins would hide this site, thus explaining the change in MW and altered maturation. To gain insight into the molecular context, AlphaFold 3 was utilized to predict an interaction between RNF13 and IDS precursor form with an Asn246 glycosylated residue. Interestingly, the predicted model showed that the interaction between RNF13 and IDS was still possible, thanks to a slight repositioning of the residues. Indeed, while the interaction between Glu245 of IDS and RNF13 remains unchanged, Asn246 no longer directly participates in the interaction when modified by the glycan chain (Fig. 4A). Instead, it is the glycan chain that now contributes to the interaction, with Arg98 and Leu100 of RNF13 (Fig 4A). To experimentally assess if RNF13 alters the glycosylation of this site, site-directed mutagenesis substituted the asparagine with a glutamine (Asn246Glu, referred to as N246Q hereafter) to inhibit glycosylation while minimizing structural alterations. Co-immunoprecipitation assays showed an increased abundance of higher MW species of N246Q variant in the immunopurified complex when compared to the WT (Fig. 4B). In the lysate, a difference in MW is observed for N246Q compared to WT when RNF13 is present (purple arrow; Fig. 4B). An additional shift for N246Q+RNF13 suggests that RNF13 likely alters another glycosylation site. Furthermore, while the mature form abundance is significantly decreased for N246Q, a difference in MW remains, contrasting with IDS WT coexpressed with RNF13 (orange arrow; Fig. 4B). No significant difference is observed between WT and N246Q for the 90kDa form, but N246Q underglycosylated abundance was higher than the WT with RNF13 (red and blue arrows, respectively; Fig. 4B-E). As the 90kDa form is produced in the TGN, it could indicate that N246Q passes through the Golgi but is not recognized by M6P receptors and that RNF13 acts upstream. IDS N246Q precursor form abundance was slightly higher than the WT (purple arrow; Fig. 4B, F). Also, the low mature abundance of the variant N246Q was decreased by half with RNF13 (orange arrow; Fig. 4B, G-H), suggesting an additive effect. Overall, our results suggest that the Asn246 glycosylation site is important for IDS maturation but is not the one affected by RNF13.

**Figure 4:**
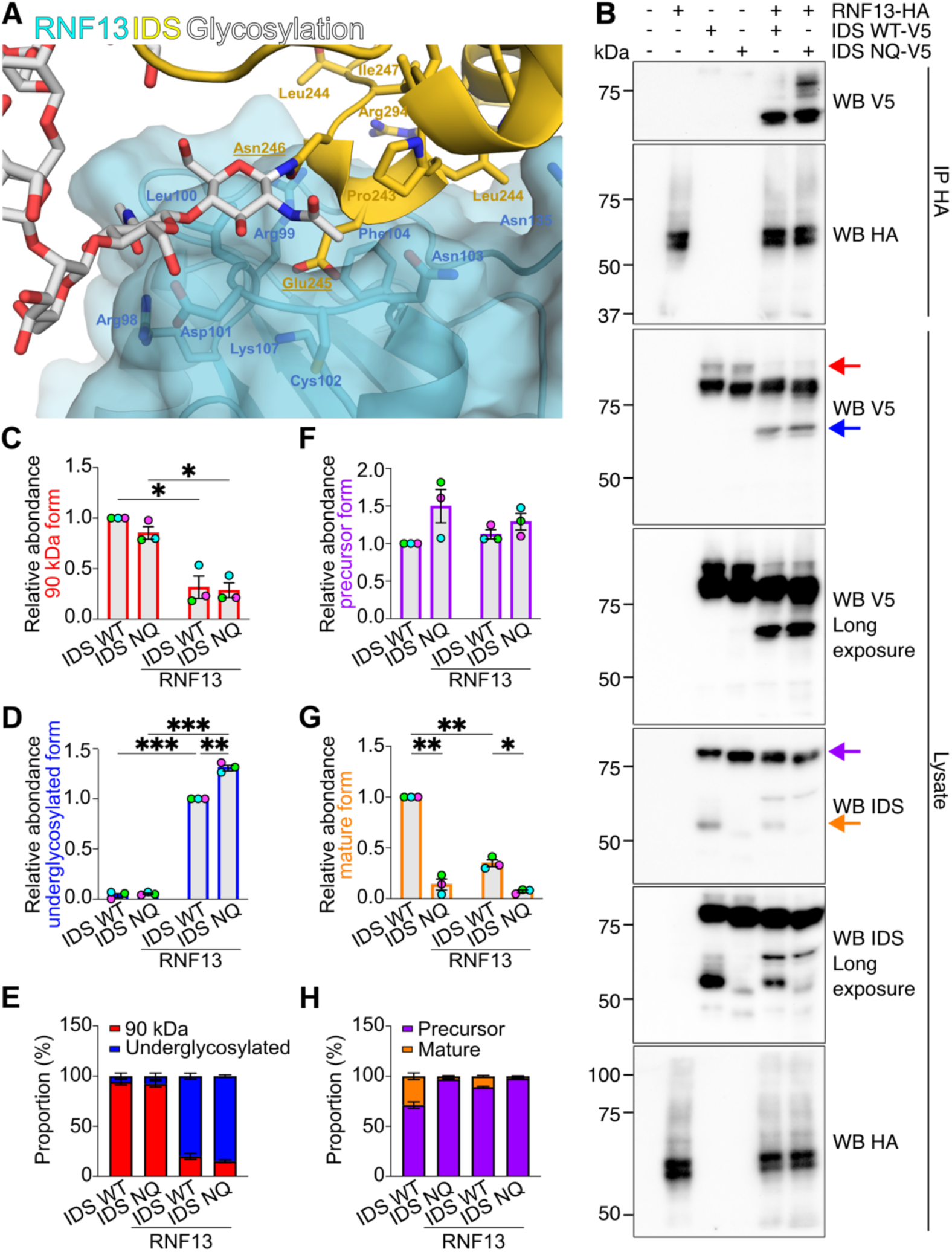
IDS glycosylation site N246 is important for its processing but is not the site affected by RNF13. (**A**) AlphaFold 3 predicted an interaction between RNF13 (cyan) and the precursor form of IDS (yellow) when the N246 is glycosylated (white). (**B**) Representative (N=3) immunoblots show IDS-V5 and RNF13-HA detection in HA-immunoprecipitated complexes from transfected HEK293T/17 cell lysates. (**C-D, F-G**) The strip plot represents the relative protein levels of IDS, specifically (**C**) 90 kDa (red arrow), (**D**) underglycosylated (blue arrow), (**F**) precursor (purple arrow) and (**G**) mature (orange arrow) forms. (**E, H**) The summary bar plots represent the proportion of (**E**) 90kDa/underglycosylated and (**H**) precursor/mature form. All results are mean ± SEM from N=3. * *p* < 0.05; ** *p* < 0.01; *** *p* < 0.001 (two-way RM-ANOVA with multiple comparisons).

To confirm if the lysosomal localization of IDS N246Q is altered, an immunofluorescence assay was performed on HEK293T/17 cells expressing RNF13, IDS WT or N246Q and Rab7. The results show that colocalization between RNF13 and N246Q is increased compared to IDS WT (Fig. 5A-B, D). A significant decrease in the MOC of IDS over Rab7 is observed for N246Q only when RNF13 is present (Fig. 5C). Additionally, RNF13/Rab7 MOC is reduced by IDS WT and by N246Q (Fig. 5E). Interaction between RNF13 and IDS in the ER could be consistent with a delay in trafficking toward the endolysosomal pathway for both proteins.

**Figure 5:**
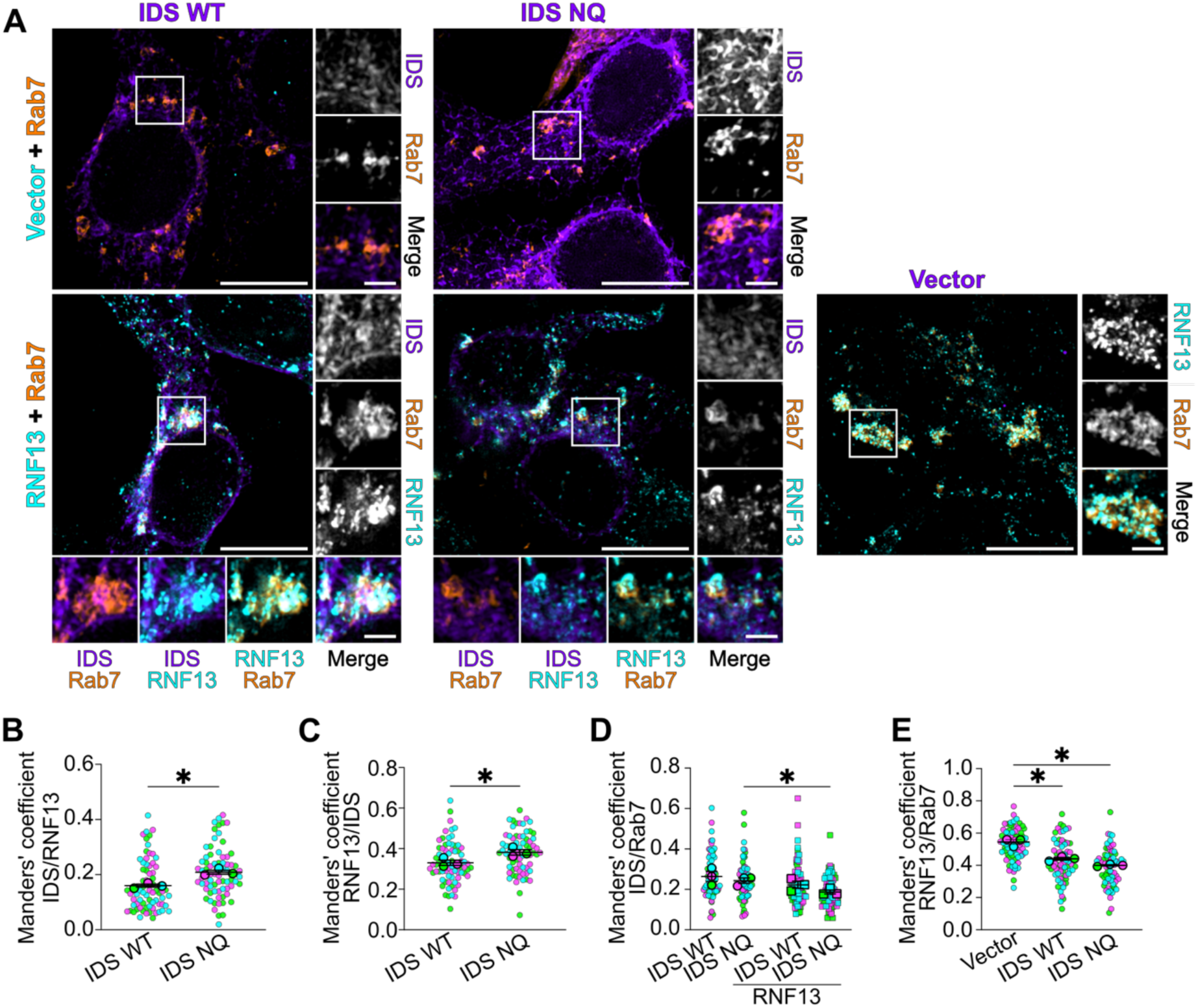
IDS N246Q has a higher colocalization with RNF13 than IDS WT. (**A**) Representative immunofluorescence images show HEK293T/17 cells expressing IDS-V5 (purple), RNF13-HA (cyan) and GFP-Rab7 (orange). Scale bars indicate 10 μm for whole cell image and 2 μm for higher magnification. (**B-E**) The jittered individual value plot represents MOC of (**B**) IDS over RNF13, (**C**) RNF13 over IDS and (**D**) IDS or (**E**) RNF13 over Rab7. All results are mean ± SEM from N=3. Larger points represent the mean of an independent experiment (N=3) whereas smaller points represent each individual cell analyzed (n=72). * *p* < 0.05; ** *p* < 0.01; *** *p* < 0.001 ((**B-C**) two-tailed paired t test or (**E**) one-or (**D**) two-way RM-ANOVA with multiple comparisons).

### RNF13 alters IDS glycosylation

To better understand how RNF13 affects IDS glycosylation, lysates of HEK293T/17 cells were treated with Endo H or Peptide:N-glycosidase F (PNGase F). Detection with anti-IDS reveals that the Endo H-treated glycosylated precursor form shifts slightly below the underglycosylated form caused by exogenous expression of RNF13 in the non-digested sample (blue arrows; Fig. 6A). The mature form also shifts but displays a smear corresponding to species resistant to Endo H that is hardly visible with RNF13 overexpression (orange arrows; Fig. 6). As Endo H-resistant glycans are added in the Golgi, the absence of smears is consistent with upstream altered trafficking. Treatment with PNGase F gives similar bands MW with or without RNF13 (Fig. 6), suggesting that the differences observed so far are caused by changes in glycosylation patterns. Importantly, the underglycosylated form caused by RNF13 is slightly higher than the one obtained with Endo H digestion but slightly lower than with PNGase F, suggesting that some residual Endo H-resistant species might be on this form (blue arrows; Fig. 6). Overexposition of the immunoblot to observe lower MW bands show that undigested IDS presents very different band patterns when RNF13 is present (Fig. 6). When treated with Endo H, the bands around 52 and 35 kDa present a small shift with RNF13 while digestion with PNGase F restablish identical bands MW (Fig. 6), suggesting that changes in Endo H-resistant glycosylation cause the shift. Because the V5-tag is on IDS C-terminus, they most likely represent cleaved C-terminal SD2 fragments [6]. The MW is higher than expected (14 kDa) but might be caused by the dimerization of the protein through its large hydrophobic region [4]. These results suggest that RNF13 mostly affect IDS glycosylation resistant to Endo H, which is consistent with an interaction occurring in the ER.

**Figure 6:**
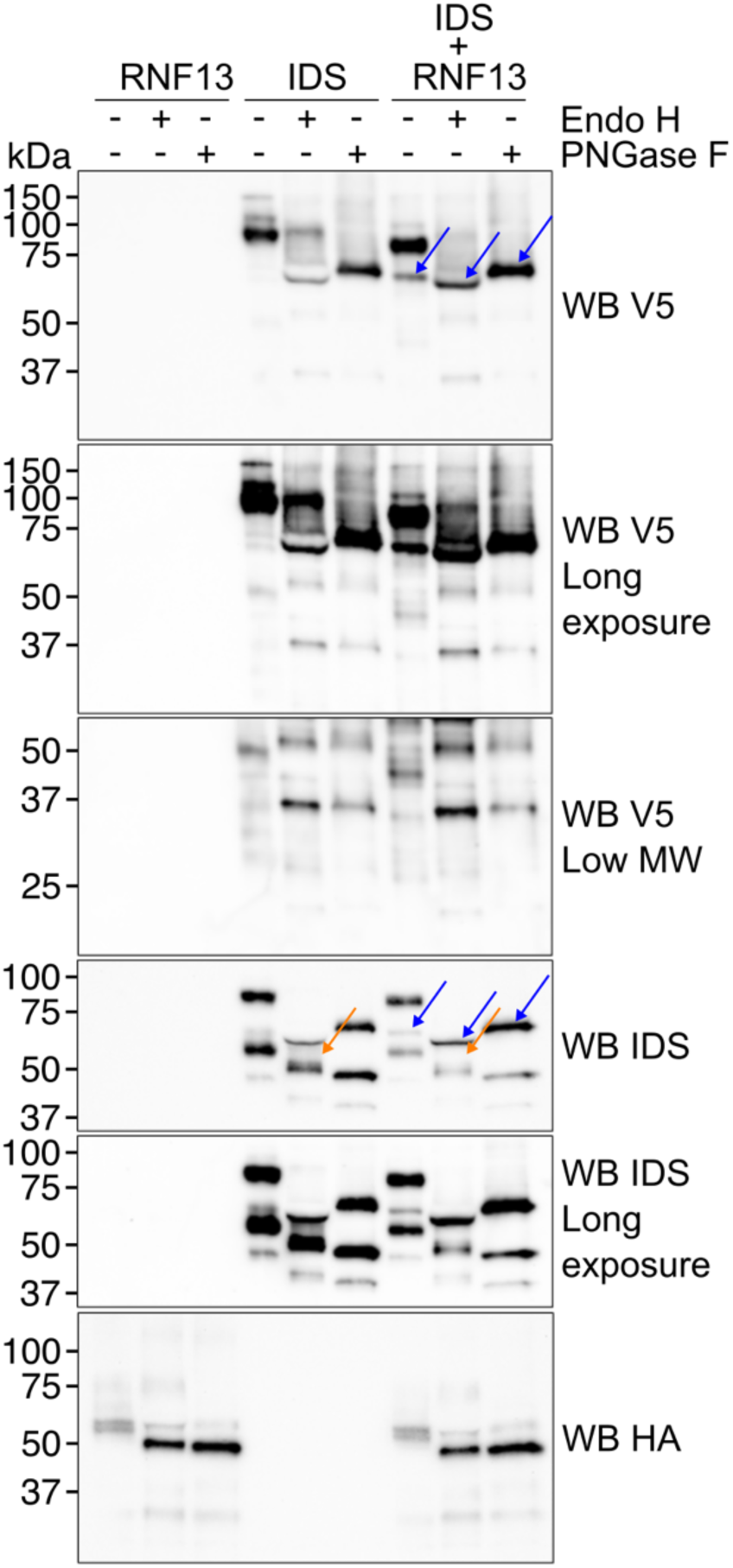
RNF13 decreases Endo H resistant species of the mature form and greatly modifies the SD2 glycosylation. Representative immunoblots show IDS-V5 and RNF13-HA detection in transfected HEK293T/17 cell lysates digested with Endo H or PNGase F (N=4). The blue arrows represent the underglycosylated form caused by RNF13, and the orange arrows represent the Endo H-resistant glycans.

### RNF13 does not affect IDS glycosylation by inducing ER stress

Knowing that ER stress can induce glycosylation change [28] and that exogenous RNF13 expression increases ER stress [14], it is possible that RNF13 indirectly impacts IDS glycosylation. To test this, HEK293T/17 cells expressing RNF13 and IDS were treated with either tunicamycin (Tn) to inhibit N-glycosylation or thapsigargin (Tg) to induce ER stress without directly affecting glycosylation. As expected, treatments resulted in an increased level of endogenous IRE1α protein when compared to the vehicule, confirming their efficacy (Fig. 7A). The IDS deglycosylated band pattern observed with Tn treatment, but not Tg, was similar to untreated IDS combined with RNF13 (blue arrow; Fig. 7B). While Tn treatment decreased the abundance of the IDS precursor form only when IDS is expressed alone, no treatment affected IDS mature form, regardless of RNF13 overexpression (purple and orange arrows, respectively; Fig. 7B-D). In contrast to IDS alone, these results suggest that the precursor form synthesized prior to the Tn treatment was unable to mature in the presence of RNF13. In addition, these results suggest that RNF13 does not affect IDS glycosylation by inducing ER stress.

**Figure 7:**
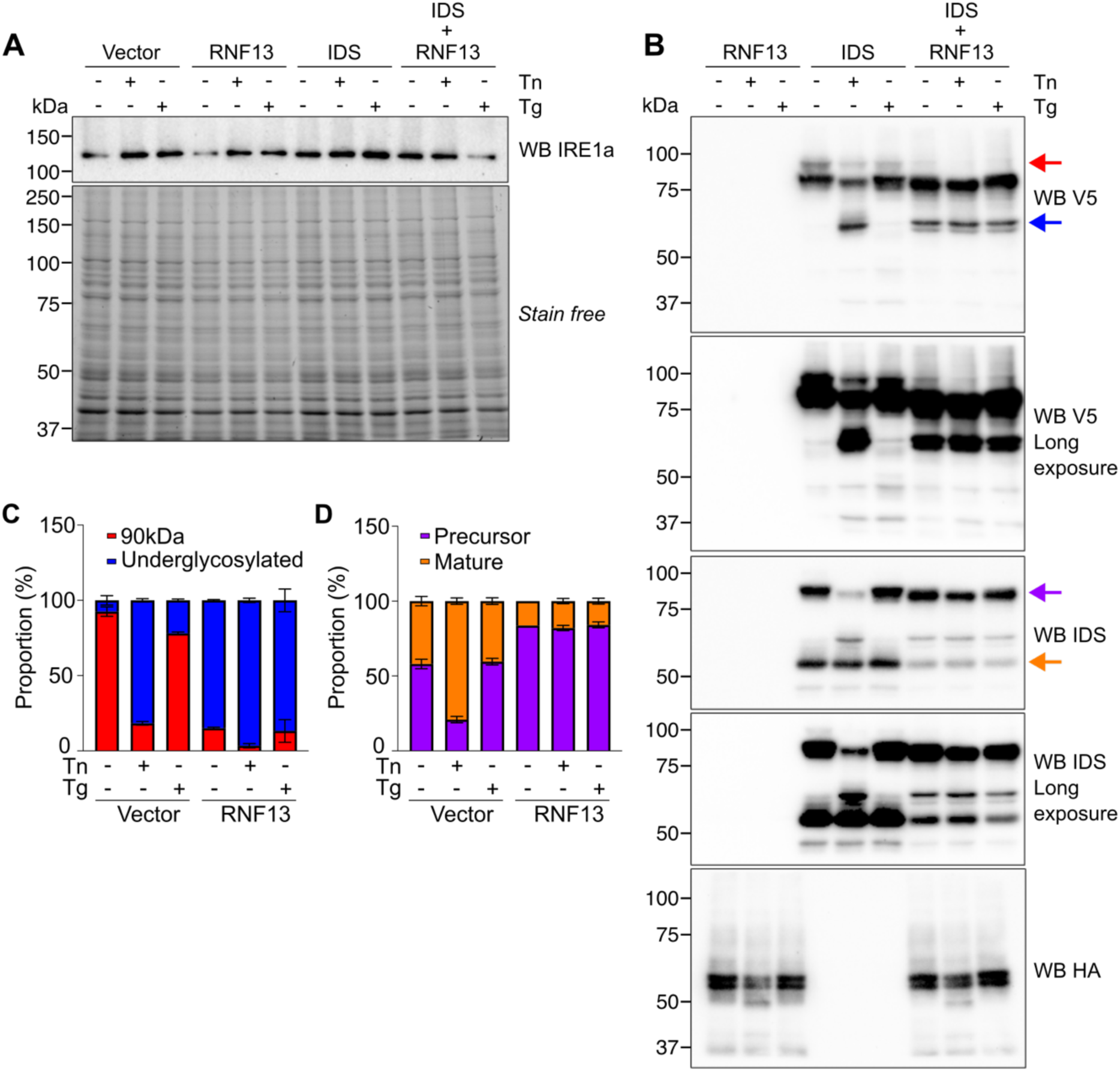
RNF13 inhibits IDS maturation but not ER stress induction. (**A-B**) Representative immunoblots show (**A**) endogenous IRE1α or (**B**) IDS-V5 and RNF13-HA detection in transfected HEK293T/17 cell lysates treated with Tn for 4 hours or Tg for 1 hour (N=3). The arrows represent the 90 kDa (red), underglycosylated (blue), precursor (purple), and mature (orange) forms. (**C-D**) The summary bar plots represent the proportion of (**C**) 90kDa/underglycosylated and (**D**) precursor/mature forms.

### RNF13/RNF167 heterodimerize and modify the interaction and abundance of IDS precursor and underglycosylated forms

While RNF13 effect on IDS is not caused by inducing ER stress, both proteins are translated and glycosylated using similar machinery in the ER. To control for the effect of machinery saturation, HEK293T/17 cells were transiently transfected with plasmids encoding the proteins IDS-V5, RNF13-HA and RNF167-YFP, another E3 ligase from the PA-TM-RING family. Interestingly, RNF167 and RNF13 had different effects on IDS, supporting that the phenotypes observed are specific and not caused by defects in ER quality control or trafficking pathways (Fig. 8A). While both the 90 kDa and mature forms of IDS were similarly affected by RNF13 and RNF167, the underglycosylated form was barely present with RNF167 (red, orange and blue arrows, respectively; Fig. 8A-D, F-G). Intriguingly, combination of RNF13 and RNF167 highly increased this form compared to RNF13 alone (blue arrow; Fig. 8A, C-D). In addition, the precursor form band pattern was different with RNF167, as it appeared less intense and like a small smear where multiple bands were almost distinguishable, but the overall abundance was not significantly different (purple arrow; Fig. 8A, E). RNF13 increased the precursor form abundance while the addition of RNF167 restored its level (purple arrow; Fig. 8A, E). Intriguingly, an additional shift of the band MW is observed when RNF13 and RNF167 are combined (purple arrow; Fig. 8A). While decreasing the mature form, RNF167 barely modified the ratio of precursor/mature form, which importantly contrasts when RNF13 is present (Fig. 8A, G). RNF167 was not affected by the different conditions, but RNF13 bands were slightly shifted in the presence of RNF167 (cyan arrow; Fig. 8A). RING finger domain-containing E3 ligases can function as monomers, dimers or multi-subunit complexes [29]. Homodimerization of RNF13 has been suggested, whereas it has been experimentally demonstrated for RNF167 [30]. In addition, RNF13 and RNF167 are identified as interactors on the BioGRID database [13, 31]. To gain insight into this possible heterodimer, AlphaFold 3 was used to predict the interaction between RNF13 and RNF167 proteins. As expected, they interact through multiple regions (Fig. 8H). Notably, both transmembrane domains, which are mostly composed of hydrophobic residues, interact with one another (Fig. 8H), although an important contribution comes from their PA domain in the N-terminal domains (Fig. 8H). Interestingly, the surface of RNF13 interacting with RNF167 is the same one as IDS. The C-terminal region, or RING domain, does not contribute to the dimerization (Fig. 8H). Immunofluorescence assays were performed in HeLa cells expressing RNF13-GFP or RNF167-HA to determine the subcellular localization of this complex by detecting the endogenous Lamp1 protein. The images show that RNF13 and RNF167 highly colocalize together, and both proteins colocalize with Lamp1 (Fig. 8I). However, presence of RNF13 significantly increased the MOC of RNF167/Lamp1 while RNF167 decreased it for RNF13/Lamp1, although not significantly (Fig. 8I-K), which could suggest that the heterodimer trafficking and/or function might be different than the monomers and/or homodimers. To better understand how the heterodimer interacts with IDS, co-immunoprecipitation assays were performed from HEK293T/17 cell lysates expressing RNF13-HA, RNF167-YFP and IDS-V5 proteins. The lysates were divided to use both anti-HA and anti-GFP beads to determine which IDS form interacts with RNF13 or RNF167, respectively. Both RNF13 and RNF167 were detected in the immunoprecipitated complex of the other ligase, confirming the heterodimerization (Fig. 8L). More specifically, in contrast to the two predominant bands detected in the lysate, RNF13 band pattern in the immunopurified complex revealed enrichment of the deglycosylated band (around 42 kDa) in addition to a heavily glycosylated form, represented by a smear above 50 kDa (WB HA in IP GFP; Fig. 8L). Interestingly, while RNF167 interacted predominantly with the precursor form of IDS and RNF13 with the underglycosylated IDS form, the complex containing RNF13/RNF167/IDS contains both forms (WB V5 in IP GFP and HA, respectively; Fig. 8L). In addition, a higher abundance of IDS is found in both immunopurified complexes when both RNF13 and RNF167 are present (WB V5 in both IP; Fig. 8L). Overall, these results suggest that RNF13/RNF167 heterodimer trafficking is regulated in a distinct manner compared to the individual proteins and that each complex (RNF13, RNF167 and heterodimer RNF13/RNF167) interacts and affects IDS differently.

**Figure 8:**
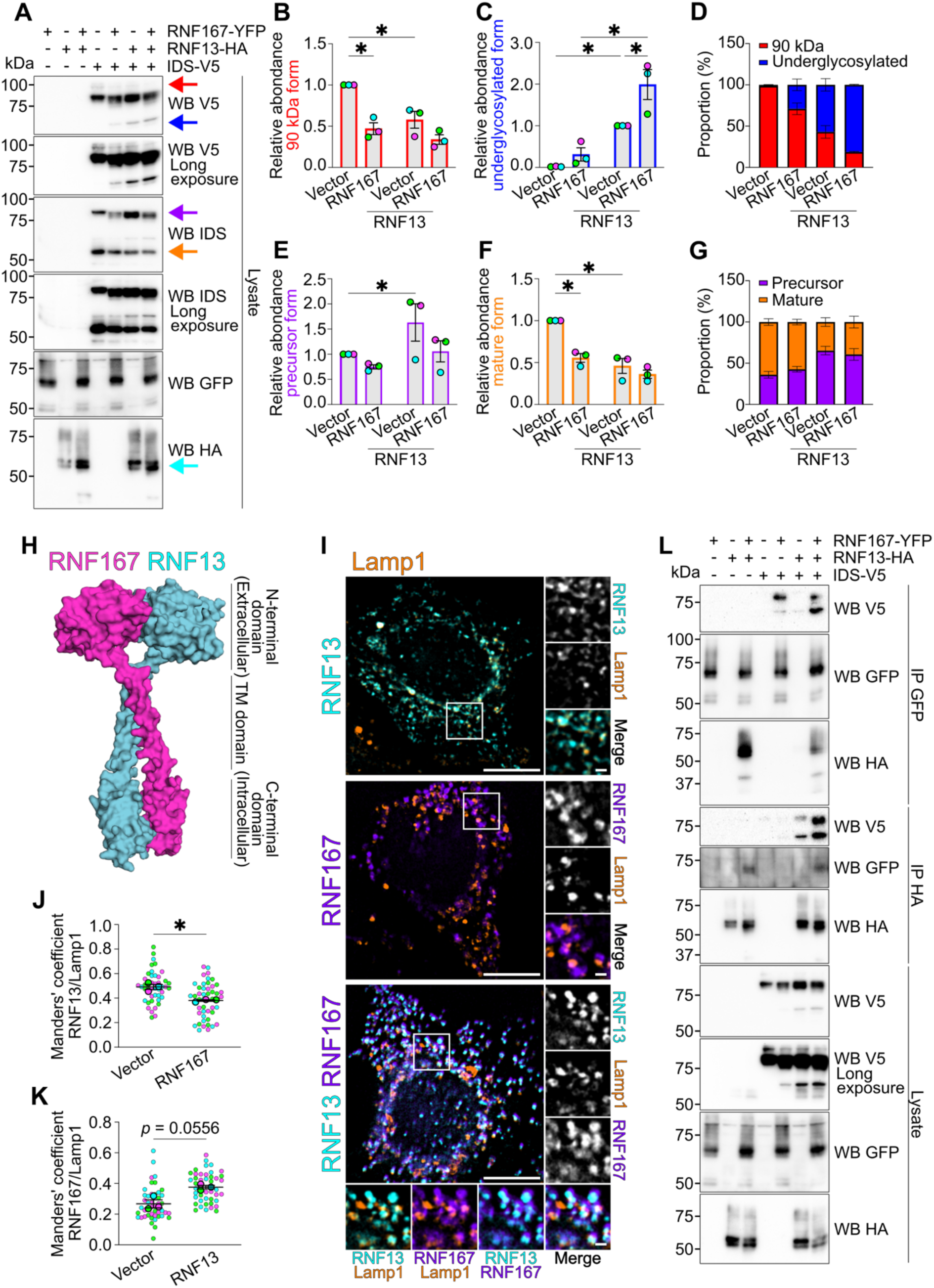
RNF13/RNF167 heterodimer modifies the interaction with IDS, increasing the abundance of the underglycosylated form. (**A**) Representative (N=3) immunoblots show IDS-V5, RNF167-YFP and RNF13-HA detection from transfected HEK293T/17 cell lysates. The arrows represent the 90kDa (red), underglycosylated (blue), precursor (purple) and mature (orange) form of IDS and RNF13 (cyan). (**B-C, E-F**) The strip plots represent relative protein levels of IDS (**B**) 90kDa, (**C**) underglycosylated, (**E**) precursor and (**F**) mature forms. (**D, G**) The summary bar plots represent the proportion of (**D**) 90kDa/underglycosylated and (**G**) precursor/mature form. (**H**) AlphaFold 3 predicted an interaction between RNF13 (cyan) and RNF167 (purple). The model is represented without the signal peptide (RNF13: a.a. 1-34; RNF167: a.a. 1-24) and the disordered C-terminal region (RNF13: a.a. 285-381; RNF167: a.a. 271-350). These regions do not affect the predicted model. (**I**) Representative immunofluorescence images show endogenous detection of Lamp1 (orange) in HeLa cells expressing RNF13 (cyan) and RNF167 (purple). Scale bars indicate 10 μm for whole cell image and 2 μm for higher magnification. (**J-K**) The jittered individual value plot represents MOC of (**J**) RNF13 or (**K**) RNF167 over Lamp1. Each color represents an independent experiment (N=3). Smaller points represent individual cells analyzed (n=45). Larger points represent the mean of an independent experiment. (**L**) Representative (N=3) immunoblots show IDS-V5, RNF167-YFP and RNF13-HA detection in immunopurified complexes with anti-HA or anti-GFP beads from transfected HEK293T/17 cell lysates. All results are mean ± SEM from N=3. * *p* < 0.05 ((**B-C, E-F**) two-way RM-ANOVA with multiple comparisons or (**J-K**) two-tailed paired t test).

### Knocking down RNF13 or RNF167 modifies how the other E3 ligase affects IDS

Thus far, this study demonstrates that RNF13 alters IDS glycosylation and maturation, and that the addition of exogenous RNF167 enhances the phenotype. However, since we have demonstrated that RNF13 and RNF167 heterodimerize, it is possible that some of the endogenous RNF13 or RNF167 heterodimerize with the overexpressed RNF167 or RNF13, respectively, and alter the phenotype that we have characterized so far. To both control if the phenotypes observed are caused by overexpression and to determine the role of the monomers and/or homodimers, HEK293T/17 cells were transfected with small interfering RNAs (siRNAs) to knockdown either RNF13 or RNF167. Whereas the efficiency of DsiRNAs against RNF167 was previously confirmed [32], two DsiRNAs against RNF13 were tested against the exogenous RNF13-HA (Fig. 9A-B). This approach was selected due to the limited success, at least in our hands, of the antibodies against endogenous RNF13. First, we determined whether the subcellular localization of RNF13 and RNF167 differed when the other ligase was repressed by detecting endogenous Lamp1 in HeLa cells expressing RNF167-HA or RNF13-HA and transfected with DsiRNAs against RNF13 or RNF167, respectively. In concordance with the previous results (Fig. 8I-K), RNF167 colocalization with Lamp1 was decreased by RNF13 knockdown while the MOC of RNF13 over Lamp1 was increased when RNF167 was repressed (Fig. 9C-F). This suggests that RNF13 is preferentially targeted to lysosomes compared to RNF167, and that the heterodimer influences their position in the endolysosomal pathway. Then, in HEK293T/17 cells expressing IDS-V5, knockdown of RNF13 and RNF167 increased the 90 kDa form, independently of the other ligase expression (Fig. 9G-J). Exogenous expression of RNF167, combined with RNF13 knockdown, increased the underglycosylated form compared to RNF167 alone, although not significantly, and a slight increase in the band MW could be observed (Fig. 9G, K). In contrast, knockdown of RNF167 resulted in a slight decrease in MW that did not affect IDS abundance (Fig. 9H, L). Interestingly, this form was also detected with longer exposure time for IDS alone with siRNF167, but not siCTL or siRNF13 (Fig. 9G-H). SD2 fragments were also detected at higher levels in siRNF167 conditions compared to RNF13 overexpression, whereas they were undetectable in IDS alone (Fig. 9H). Notably, the precursor form was unaffected by RNF167 overexpression or RNF13 repression (Fig. 9G, M). In contrast, RNF167 knockdown or RNF13 overexpression both increased the abundance of this form (Fig. 9H, N). Overall, these results suggest that both RNF13 and RNF167 modify IDS glycosylation and maturation.

**Figure 9:**
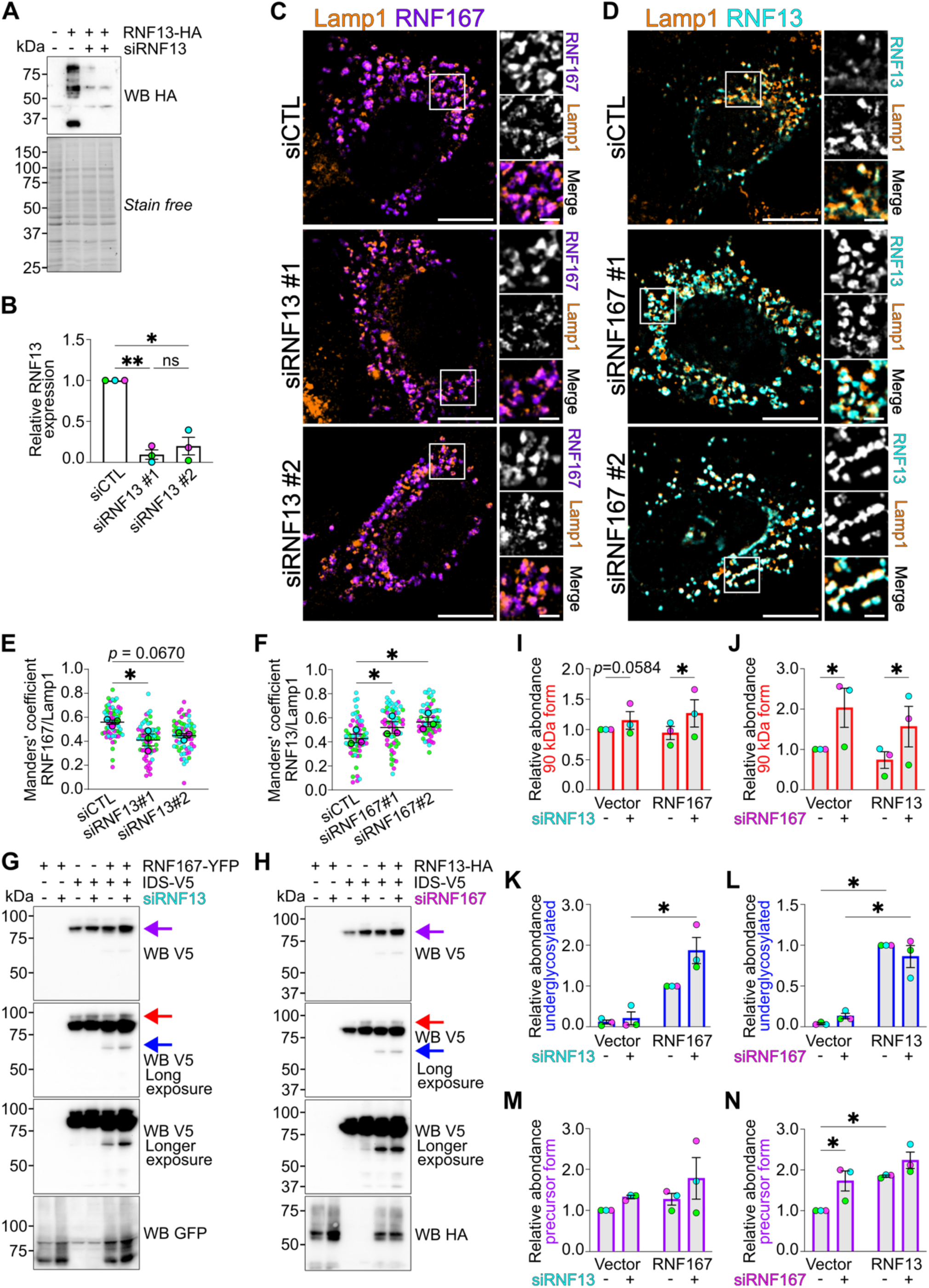
Knockdown of endogenous RNF13 or RNF167 modifies how the other E3 ligase affects IDS. (**A**) Representative (N=3) immunoblots show RNF13-HA detection from the lysate of HEK293T/17 cells transfected with siRNF13. (**B**) The strip plot represents relative protein levels of HA detected. (**C-D**) Representative immunofluorescence images show endogenous detection of Lamp1 (orange) in HeLa cells expressing (**C**) RNF167 (purple) and siRNF13 or (**D**) RNF13 (cyan) and siRNF167. Scale bars indicate 10 μm for whole cell image and 2 μm for higher magnification. (**E-F**) The jittered individual value plots represent MOC of (**E**) RNF167 or (**F**) RNF13 over Lamp1. Each color represents an independent experiment (N=3). Smaller points represent individual cells analyzed (n=63). Bigger points represent the mean of an independent experiment. (**G-H**) Representative immunoblots (N=3) show IDS-V5 and (**G**) RNF167-YFP or (**H**) RNF13-HA detection from transfected HEK293T/17 cell lysates with (**G**) siRNF13 or (**H**) siRNF167. (**I-N**) The strip plots represent relative protein levels of IDS (**I-J**) 90kDa (red arrow), (**K-L**) underglycosylated (blue arrow), (**M-N**) precursor (purple arrow) form. All results are mean ± SEM from N=3. * *p* < 0.05; ** *p* < 0.01 ((**B, E-F**) one-way or (**I-N**) two-way RM-ANOVA with multiple comparisons).

### RNF13 E3 ligase activity is required for a proper interaction with IDS and to increase the underglycosylated form

Generally, in RING finger protein heterodimers, one of the monomers lacks catalytic activity and stabilizes the active E2-binding RING domain of the other ligase [29]. To test if the E3 activity ligase of both RNF13 and RNF167 is needed to modulate IDS, dominant negative (DN) mutants of Ub ligase were expressed in HEK293T/17 cells with IDS-V5 and either WT or DN of the other ligase. Both RNF13 and RNF167 DN, alone or together, increased the 90 kDa form of IDS (red arrow; Fig. 10A). However, at least one of the two WT ligase protein was sufficient to restore the level of this form to baseline (red arrow; Fig. 10A). RNF167 E3 ligase activity was needed to separate the precursor form into multiple bands, independently of RNF13, although RNF13 WT was needed to cause a MW shift (purple arrow; Fig. 10A). Combined RNF13 and RNF167 increased IDS underglycosylated form compared to RNF13 alone, while this form is barely detectable with RNF167 alone (blue arrow; Fig. 10A). Expression of RNF13 DN with WT RNF167 results in IDS level similar to what was obtained with RNF167 alone (blue arrow; Fig. 10A). In contrast, RNF167 DN and WT RNF13 increased IDS underglycosylated form as much as the combination of both WT ligases (blue arrow; Fig. 10A). These results suggest that the E3 ligase activity of RNF13, but not of RNF167, is needed to increase IDS underglycosylated form.

**Figure 10:**
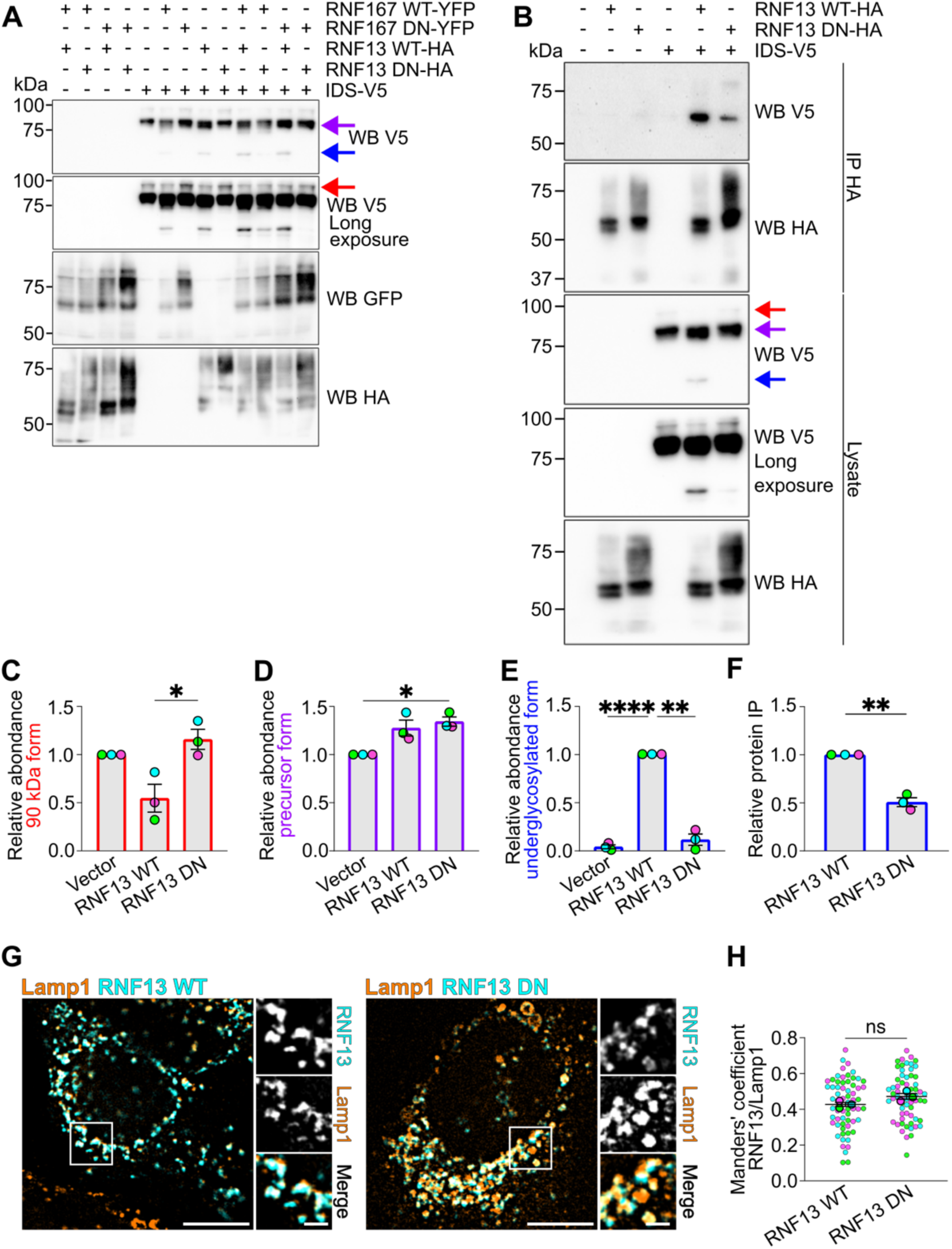
RNF13 E3 ligase activity is required to increase the underglycosylated form but not RNF167 and mutation in the RING domain of RNF13 alters the interaction with IDS. (**A**) Representative immunoblots show V5-tagged IDS WT, YFP-tagged RNF167 WT or DN and HA-tagged RNF13 WT or DN detection from transfected HEK293T/17 cell lysates (N=2). (**B**) Representative (N=3) immunoblots show IDS-V5 and RNF13 WT or DN-HA detection in HA-immunoprecipitated complexes from transfected HEK293T/17 cell lysates. (**C-E**) The strip plots represent relative protein levels of IDS (**C**) 90kDa (red arrow), (**D**) precursor (purple arrow) and (**E**) underglycosylated (blue arrow) forms. (**F**) The strip plot represents the normalized amount of V5 detected in the immunoprecipitated complex over the amount in the total lysate and over the quantity of RNF13 that was immunoprecipitated. (**G**) Representative (N=3) immunofluorescence images show endogenous detection of Lamp1 (orange) in HeLa cells expressing RNF13-HA (cyan). Scale bars indicate 10 μm for whole cell image and 2 μm for higher magnification. (**H**) The jittered individual value plot represents MOC of RNF13 over Lamp1. Each color represents an independent experiment. Smaller points represent individual cells analyzed (n=63). Bigger points represent the mean of an independent experiment. All results are mean ± SEM from N=3. ns = not significant, * *p* < 0.05; ** *p* < 0.01; **** *p* < 0.0001 ((**C-E**) one-way RM-ANOVA with multiple comparisons or (**F, H**) two-tailed paired t test).

RNF13 RING domain is localized in the cytosol, while RNF13’s PA domain and IDS are localized in the lumen [22] and predicted to associate together (Fig. 2A). However, RNF13 DN display a different glycosylation pattern than RNF13 WT (Fig. 10A). Because the mutations in the RING domain are on the cytosolic side, we wondered if IDS/RNF13 interaction could be affected. To test this, anti-HA co-immunoprecipitations were performed from HEK293T/17 cell lysates expressing either wild-type (WT) or dominant-negative (DN) RNF13-HA with IDS-V5 proteins. RNF13 DN increased IDS 90 kDa when compared to RNF13 WT (red arrow; Fig. 10B-C). Additionally, RNF13 DN increased significantly the precursor form, slightly more than RNF13 WT (purple arrow; Fig. 10B, D). As obtained previously, the IDS underglycosylated form was not detectable with RNF13 DN in the lysate; however, a small quantity was detected in the immunopurified complex (blue arrow; Fig. 10B, E). To account for the fact that the underglycosylated form was barely detectable in the lysate but that RNF13 DN expression is higher, we normalized the amount of V5-tagged protein detected in the immunoprecipitated complex relative to the total lysate and the amount of RNF13 immunoprecipitated (Fig. 10 B, F). As a result, RNF13 DN immunoprecipitated approximately half the amount of IDS compared to WT RNF13, suggesting that the RING domain activity modulates the interaction, but is not absolutely required (Fig. 10B, F). To ensure that the interaction of DN RNF13 with IDS is not an artifact of a prolonged ER retention caused by trafficking defects, immunofluorescence assays were performed in HeLa cells expressing RNF13 WT or DN. The results show that both proteins colocalize with Lamp1-positive lysosomes, and that the MOC of RNF13 DN/Lamp1 is not significantly different than the WT (Fig. 10G-H). Overall, these results suggest that the E3 ligase activity of RNF13 is required to induce the underglycosylated form of IDS, but not for the mutual interaction.

### RNF13 protects IDS underglycosylated form and its C-terminal fragment from rapid proteasomal degradation

As glycosylation trimming targets glycoproteins for ER-associated degradation (ERAD) [33] and the effect of RNF13 on the underglycosylated form of IDS is dependent on its E3 ligase activity, we wondered if the altered IDS maturation caused by RNF13 is delayed or promotes its degradation. To test this, HEK293T/17 cells expressing RNF13 and IDS were treated with CHX for 30 minutes before either the lysosomal degradation inhibitor chloroquine (CQ) or the proteasome inhibitor MG132 was added for 4 hours. RNF13 abundance increased with both treatments [16, 34], confirming their efficacy (Fig. 11A, H). Both the 90kDa and underglycosylated forms were decreased by CHX independently of RNF13, whereas treatment of CQ increased the abundance of the 90 kDa form (red and blue arrows, respectively; Fig. 11B-D). In contrast, CQ treatment decreased IDS mature form, reducing its proportion by 10% (orange arrow; Fig. 11F-G). While the CHX was very efficient to decrease the precursor form of IDS alone, this effect was inhibited by RNF13 presence (purple arrow; Fig. 11E). Furthermore, the low MW bands detected by anti-V5 but not by anti-IDS, indicating that they correspond to C-terminal SD2 fragments, were decreased by CQ only when RNF13 was present (Fig. 11A). Independently of the treatments, RNF13 increased the IDS C-terminal fragments abundance when compared to IDS alone and most bands have different MW, further supporting the glycosylation change (Fig. 11A). Overall, these results suggest that IDS is not rapidly degraded by the lysosomes.

**Figure 11:**
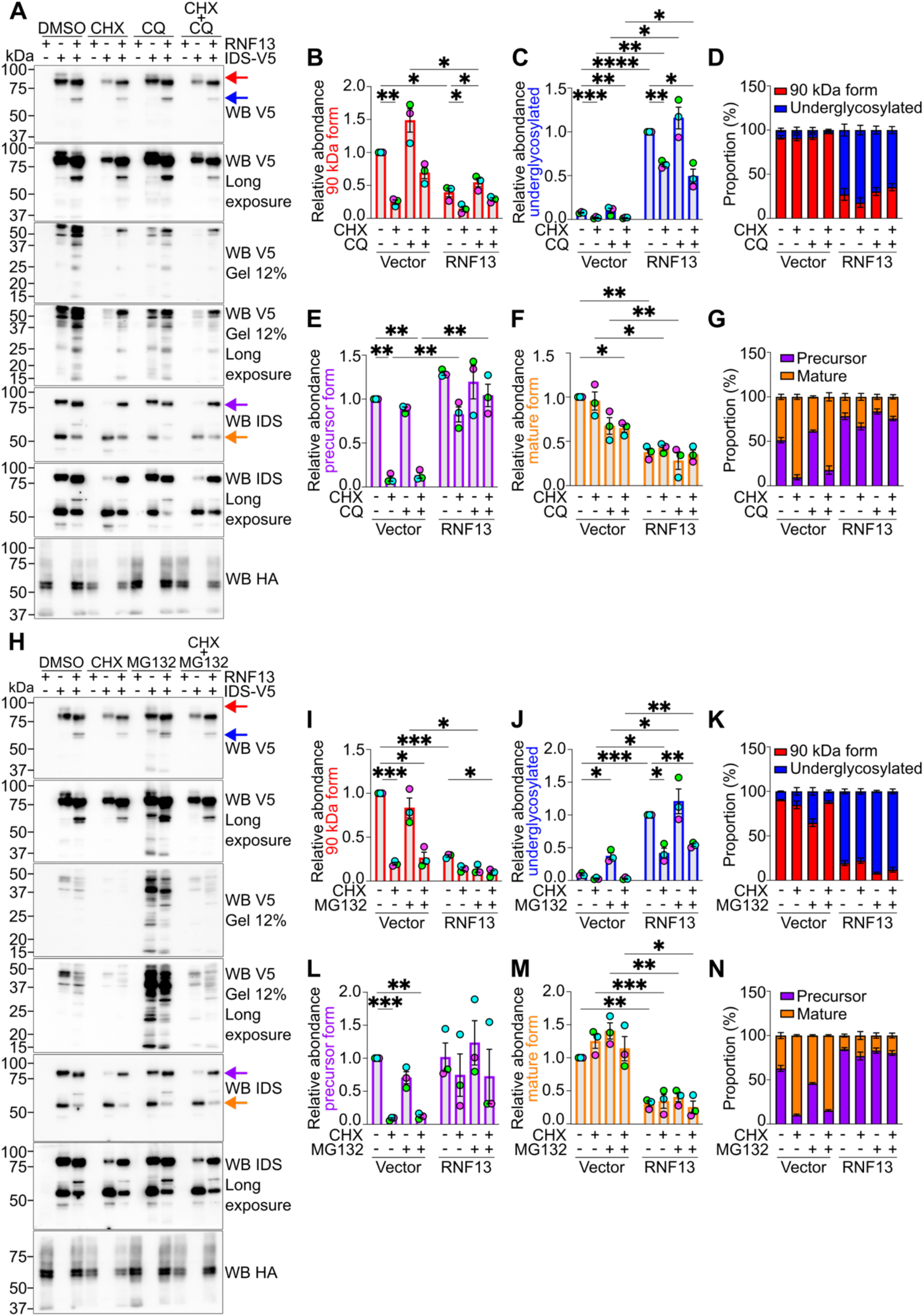
IDS is not degraded by the lysosomes but by the proteasome while RNF13 protects some forms from degradation. (**A, H**) Representative (N=3) immunoblots show IDS-V5 and RNF13-HA detection in lysates from transfected HEK293T/17 cells treated with CHX for 30 minutes before adding (**A**) CQ or (**H**) MG132 for 4 hours. (**B-C, E-F, I-J, L-M**) The strip plots represent relative protein levels of IDS (**B, I**) 90kDa (red arrow), (**C, J**) underglycosylated (blue arrow), (**E, L**) precursor (purple arrow) and (**F, M**) mature (orange arrow) forms. (**D, G, K, N**) The summary bar plots represent the proportion of (**D, K**) 90kDa/underglycosylated and (**G, N**) precursor/mature forms. Values are mean ± SEM from N=3. Each color in the strip plots represents an independent experiment. * *p* < 0.05; ** *p* < 0.01; *** *p* < 0.001; **** *p* < 0.0001 (two-way RM-ANOVA with multiple comparisons).

The underglycosylated form of IDS increases with MG132 (blue arrow; Fig. 11H, J). Specifically, the levels of IDS alone were now comparable to the levels obtained in the presence of RNF13 treated with CHX with and without MG132 (blue arrow; Fig. 11H, J). Furthermore, MG132 increased the mature/precursor ratio for IDS but not when RNF13 was overexpressed (purple and orange arrows; Fig. 11L-N). MG132 led to an important accumulation of low MW bands detected by anti-V5 (Fig. 11H). Importantly, the band profile differed significantly with RNF13 (Fig. 11H), consistent with an effect on IDS C-terminal glycosylation pattern (Fig. 6A). It suggests that, unlike previously thought, processed IDS fragments may not be degraded by lysosomes but rather by the proteasome. These results suggest that the proteasome rapidly degrades IDS in its underglycosylated form, and that RNF13 delays the proteasomal degradation of IDS underglycosylated precursor and the SD2 fragments.

## DISCUSSION

This work presents a novel interaction between RNF13 and IDS. RNF13 appears to have a dual functional role, reducing IDS maturation while protecting the underglycosylated form and SD2 fragments from proteasomal degradation. While we have not confirmed that RNF13’s PA domain is indeed involved in the interaction, an IDS C-terminal fragment and a predominantly underglycosylated form were co-immunoprecipitated with RNF13, concurring with both predicted AlphaFold 3 models (Fig. 1). RNF13 could either act by blocking the addition or enhancing the removal or trimming of glycosylation. As most of the N-glycosylation is added by a cotranslational process [35], and the IDS form obtained in the presence of RNF13 was similar but not exactly the same as the one obtained following Tn treatment (Fig. 7), it is plausible that RNF13 enhance the removal of glycosylation. However, there are some cases where N-glycosylation is added posttranslationally, including when the acceptor sites are located in the last 50 residues of the C-terminus [35]. As IDS has two glycosylation sites within its last 50 residues (Fig. 1), RNF13 might inhibit the addition of glycosylation on one of these sites. This would be consistent with the fact that the processed SD2 fragments have different glycosylation patterns with RNF13 (Fig. 6). Alternatively, N-glycosylation removal is generally performed by PNGase on the cytosolic side, after the retrotranslocation of a protein for its proteasomal degradation [5]. However, while plausible since our results demonstrate that the proteasome rapidly degrades this form, the band obtained by RNF13 was not exactly at the same MW than when digested with PNGase F, suggesting some residual glycosylation (Fig. 6). Instead, an interesting mechanism may exist in which the interaction RNF13/IDS keeps IDS into the ER lumen, extending IDS exposure to ER degradation-enhancing α-mannosidase-like protein (EDEM), leading to enhance mannose triming [36]. This is consistent with the observation that the reduced lysosomal targeting of the IDS variant N246Q resulted in an increased IDS underglycosylated form when RNF13 was present (Fig. 4). Moreover, RNF13 seems to protect this underglycosylated form from proteasomal degradation, as this proposed mechanism would inhibit retrotranslocation (Fig. 11).

A notable finding of our study is that the SD2 fragments are rapidly degraded by the proteasome but not by the lysosomes, suggesting that they should be retrotranslocated into the cytoplasm. While it was initially reported that IDS is processed in the lysosomes [37], a more recent study suggests that it is instead cleaved in the Golgi [7]. The Endosome and Golgi-associated degradation (EGAD) pathway was recently discovered in *Saccharomyces cerevisiae* [38]. EGAD has remarkable similarities to ERAD, as it utilizes the proteasome to degrade substrates in post-ER organelles [39]. While the substrate recognition for EGAD remains to be elucidated, it has been proposed that glycosylation patterns may play a mechanistic role [38]. As IDS glycosylation trimming is involved in its proteasomal degradation in the ER [40], SD2 fragments could be degraded by EGAD, while the RNF13-induced change in glycosylation could be protective. Importantly, as our experiments were performed using overexpression, it cannot be totally ruled out that the proteasomal machinery could be saturated. However, this hypothesis appears unlikely as the assay combining CHX and MG132 treatments demonstrate that half the underglycosylated form induced by RNF13 is degraded within the first 30 minutes while no difference is seen in the following 4 hours (Fig. 11). If the machinery was saturated, the CHX treatment alone should have significantly decrease this form to levels lower than the CHX+MG132 condition. The additional 4 hours should have allowed the proteasome to eliminate the accumulation effectively (Fig. 11). Additionally, it is important to note that the CQ treatment was performed for 4 hours, and we cannot exclude the possibility that the forms that reach the lysosomes have a longer half-life. In any case, our results suggest that processed forms of IDS can be degraded by the proteasome before reaching the lysosomes, highlighting the importance of further studying IDS trafficking and processing.

While interesting, other mechanisms may be involved in the generation of IDS underglycosylated form by RNF13. Notably, RNF13 DN interacts with approximately 50% the amount of IDS interacting with the WT while only increasing the underglycosylated form by 15%, suggesting that the interaction is not as important as RNF13 E3 ligase activity (Fig. 10). Additionally, our study reveals the existence of the RNF13/RNF167 heterodimer, where only the E3 ligase activity of RNF13 combined with the expression of RNF167 is needed to increase the level of IDS underglycosylated form (Fig. 10). This finding suggests that RNF167 may help stabilize the interaction between E2 and RNF13, similar to what has been previously described for other heterodimers in the RING protein family [29]. It could also explain why this form was slightly decreased when RNF13 and siRNF167 were combined (Fig. 9). In this case, ubiquitination of a substrate might be a plausible explanation for the expression of IDS in an underglycosylated form. As ubiquitin K48 chains are responsible for proteasomal degradation, it is possible that RNF13 ubiquitinates a different type of chain on IDS once it has retrotranslocated to the cytosol, removing its signal for degradation. However, RNF13 is known to regulate indirectly some substrates by targeting the E3 ligase responsible for their degradation [16, 41]. For example, RNF13 stabilizes TRIM29 through K63-linked ubiquitination, which in turn promotes K48-linked ubiquitination of STING leading to its degradation [16]. While speculative, it is possible that RNF13 regulates another E3 ligase, such as HRD1 [42], involved in IDS degradation. Involvement of another E3 ligase is consistent with the increased 90 kDa form by both RNF13 and RNF167 repression and DN mutants, as this form should not be retrotranslocated into the cytoplasm (Fig 9-10). As it is beyond the scope of this manuscript, further studies will be necessary to investigate whether RNF13 modifies IDS ubiquitination either directly or indirectly and how IDS ubiquitination impacts its trafficking and processing.

The SD2 fragments were also increased when RNF167 was repressed. Specifically, an increased underglycosylated form of IDS results from the following conditions: (1) comparing RNF167/RNF13 exogenous to RNF13 exogenous suggests a preference for the formation of RNF13/RNF167 heterodimer over RNF13 monomer or homodimer; (2) comparing exogenous RNF167 with siRNF13 to siCTL benefit the formation of exogenous RNF167 monomer or homodimer; and (3) comparing IDS alone with siCTL to siRNF167 led to the formation of endogenous RNF13 monomer or homodimer (Fig. 9). Based on our immunofluorescence assays with both overexpression and repression, RNF13-containing dimers exhibit higher colocalization with Lamp1 than the RNF167-containing one (Fig. 8-9). Importantly, while it is assumed that there are only three possible states (monomer, homodimer or heterodimer), there might be other members of the PA-TM-RING family that could also heterodimerize with either RNF13 and/or RNF167 and thus further change the intracellular dynamics of IDS. The involvement of another member, to be identified, could explain why both exogenous RNF167 monomer and homodimer, as well as endogenous RNF13 monomer and homodimer, increase the underglycosylated form. In addition, while the interaction is predicted to occur around Asn246, RNF13 does not seem to affect this glycosylation site. However, the precursor levels of N246Q co-immunoprecipitated with RNF13 were higher than WT IDS, which was barely detectable (Fig. 4). Interestingly, the same form was co-immunoprecipitated when RNF167 was present, whereas RNF13 species were different than the one predominant in the lysate (Fig. 8). Although the predicted model suggested that glycosylation of IDS Asn246 could help stabilize the interaction between IDS/RNF13, contrasting with the experimental results (Fig. 4), other post-translational modifications, such as RNF13 glycosylation, were not taken into account and might be important to regulate this interaction. However, we cannot exclude any beneficial effect from the glutamine used for IDS mutagenesis. Instead of Asn246, RNF13 appears to impact the C-terminal glycosylation, which is observed in both the precursor and SD2 processed fragments. The IgG-IDS fusion protein has been shown to dimerize [43], which is consistent with the abundance of hydrophobic interactions between adjacent SD2 domains in the crystal lattice of IDS [4]. The interaction appears to depend on the cellular context, as demonstrated by the changes in IDS form immunoprecipitated by RNF13, RNF167 or RNF13/RNF167 (Fig. 8). Future studies will be needed to elucidate the molecular mechanisms underlying IDS glycosylation and maturation.

In the last few years, RNF13 has gained attention, and multiple new functions have been reported for its cytosolic RING domain [15-17, 41, 44, 45]. Whereas the RNF13 PA domain remains poorly described, it has been shown to be important for its lysosomal targeting [30]. More recently, a study published on bioRxiv suggests that RNF13’s N-terminal is the primary signal for lysosomal localization [44]. However, this contrasts with our previous work that showed that RNF13 proper lysosomal localization relies on its C-terminal for a crucial interaction with AP-3 [18]. Here, we show that IDS or RNF167 expression could affect RNF13’s trafficking (Fig. 3, 8-9). As both proteins most likely interact with the PA domain, it is consistent with the N-terminal being just as important as the C-terminal for RNF13 proper localization. Interestingly, while some heterodimeric RINGs, such as BRCA1/BARD1 and Mdm2/MdmX, where the BARD1 and MdmX monomers are inactive [46], RNF13 and RNF167 appear to both be active on their own. Accordingly, the results combining both DN suggest that the E3 ligase activity of RNF13 is sufficient for inducing IDS underglycosylated form, therefore suggesting that RNF167 would act as an “inactive” partner (Fig. 10). By using *in vitro* assays to eliminate the possibility of RNF167 dimerizing with RNF13, we have previously revealed that RNF167 possesses intrinsic E3 ligase activity [47]. Both the *Drosophila melanogaster* protein Godzilla and the *Caenorhabditis elegans* protein RING-type domain-containing protein are orthologs of human RNF13, RNF167 and ZNRF4 [22]. An interesting hypothesis is that these proteins evolved separately to regulate their spatial and temporal activities, which aligns with our immunofluorescence assays on RNF13 and RNF167 (Fig. 8-9). While the C-terminal of RNF13 is important for AP-binding [19], it is possible that interaction with RNF167 through its PA domain determines whether it binds to AP-1 or AP-3, thereby modulating its trafficking pathway. This interaction could enable the cell to regulate the localization of RNF13, which is involved in multiple cellular pathways as needed [15-17, 22, 41]. For example, when IDS was overexpressed, RNF13 trafficking was delayed (Fig. 3).

A brain-specific interactome in mice has identified 187 putative partner proteins of IDS [48]. This study suggested that IDS might be involved in multiple cellular processes, including cell growth and intracellular trafficking. The present study identified RNF13 as a novel interaction partner of IDS, suggesting mutual importance for each other’s lysosomal targeting. The dileucine motif in RNF13 has been recently demonstrated to be critical for its lysosomal targeting [18, 19]. A hot spot of genetic variants surrounding the RNF13 dileucine motif [21] has been associated with severe infantile neurodevelopmental disease, for which the symptoms overlap with MPS II and with many other lysosomal storage diseases. Intriguingly, RNF13 is highly expressed in the organs most affected by Hunter syndrome, such as the respiratory system, gastrointestinal tract, spleen, liver, heart, and brain [49]. Considering the complexity of the dynamic interaction between IDS and RNF13, it is possible that multiple cellular pathways could be altered by mutations in one or the other protein in specific tissues. Importantly, the Hunter syndrome phenotype may be a lot more complex than just GAG accumulation in the lysosomes, and our findings underscore the need to explore new research avenues.

## MATERIALS AND METHODS

### Antibodies

The following antibodies were used: purified anti-HA.11 rabbit polyclonal (WB and IF: 1:1,000, BioLegend Cat# 902302, RRID:AB_2565018), purified mouse monoclonal anti-HA.11 (IF: 1:1,000, BioLegend Cat# 901503, RRID:AB_2565007), rabbit anti-GFP (WB: 1:1000, ThermoFisher Scientific, Mississauga, ON, Canada, Cat# A6455, RRID:AB_221570), rabbit monoclonal anti-V5 (WB and IF: 1:1,000, Cell Signaling Technology Cat# 13202, RRID:AB_2687461), mouse monoclonal anti-V5 (WB: 1:5,000, Invitrogen Cat# R960-25, RRID:AB_2556564), mouse monoclonal anti-IDS (WB and IF: 1:1,000, Thermo Fisher Scientific Cat# MA5-25855, RRID:AB_2723262), rabbit monoclonal anti-IRE1α (WB: 1:500, Cell Signaling Technology Cat# 3294, RRID:AB_823545), rabbit anti-Lamp1 (IF: 1:500, New England Biolabs Ltd., Cat# 9091, RRID:AB_2687579), AlexaFluor Plus 405-conjugated goat anti-rabbit (1:250, Thermo Fisher Scientific Cat# A48254, RRID:AB_2890548), AlexaFluor Plus 647-conjugated goat anti-mouse (1:1,000, Thermo Fisher Scientific Cat# A32728, RRID:AB_2633277), horseradish peroxidase (HRP)-conjugated goat anti-mouse (1:10,000, Cell Signaling Technology Cat# 7076, RRID:AB_330924) and HRP-conjugated goat anti-rabbit (1:10,000, Cell Signaling Technology Cat# 7074, RRID:AB_2099233).

### Molecular Biology

As previously described [19], the wild-type, full-length human RNF13 transcript variant 1 (accession NM_007282.4) cDNA was synthesized (GenScript, Piscataway, NJ, USA) in pUC57 vector then excised and cloned into the pCAGGS-IRES-mCherry vector. HA- and EYFP-tagged human RNF167 were previously described [50]. For the immunofluorescence assays with RNF167 and Lamp1, the pUC57 vector was used as a template to generate an optimized Kozak (AACGAG) to the insert encoding the RNF13-GFP fusion protein. Wild-type, full-length human IDS transcript variant 1 (accession NM_000202.8) cDNA was synthesized (GenScript, Piscataway, NJ, USA) and then cloned into pcDNA3.1(+) using HindIII and XhoI. This construct served as template to generate protein variant N246Q via site-directed mutagenesis. The plasmids were confirmed by Sanger sequencing (Genome Quebec, Montreal, QC, Canada).

### Cell Culture and Transfection

HEK293T/17 (American Type Culture Collection, Gaithersburg, Cat# CRL-11268) and HeLa (from Dr. Diana Alison Averill, Montreal) cells were cultured at 37°C with a humidified atmosphere containing 5% CO_2_. No antibiotics were used for maintaining the cell lines in Dulbecco’s Modified Eagle Medium (DMEM, Thermo Fisher Scientific, Cat# 11995-065) supplemented with 10% fetal bovine serum (FBS, Wisent, Cat# 090150). Cells were not recently authenticated nor tested for mycoplasma contamination. For biochemical assays, 6-well plates were treated with 0.1 mg/mL poly-D-lysine (Sigma-Aldrich, Cat# P7280) for at least 30 minutes at room temperature. For immunofluorescence assays, 24-well plates containing a 12 mm round glass coverslip #1.5 (UltiDent Scientific Inc., St-Laurent, QC, Canada, Cat# 170-C12MM) were treated with poly-D-lysine overnight at 37°C. HEK293T/17 cells were transiently transfected using Lipofectamine 2000 transfection reagent as previously described [51]. For 24-well plates, all the volume and quantity were scaled down using a factor of 5 from 6-well plates. Transfected cells were maintained at 37°C with 5% CO_2_ for 48 hours before lysis or fixation. HeLa cells were transfected with a calcium phosphate-DNA precipitate as previously described [19], and incubated for 24 hours. When required, to induce ER stress, cells were incubated at 37°C with 5% CO_2_ with 1 µg/mL tunicamycin (Cayman Chemical, Cat #11445) or DMSO for 3 hours before adding 0.5 µM thapsigargin (Cayman Chemincal, Cat #10522) or DMSO for 1 hour. To inhibit protein synthesis, cells were incubated at 37°C with 5% CO_2_ with 25 µg/mL cycloheximide (Sigma-Aldrich, Cat# C1988) or DMSO for 30 minutes before adding either 20 µM MG132 (UBPBio, Cat# F1101) or DMSO to inhibit proteasomal degradation or 50 µg/mL chloroquine (Sigma-Aldrich, Cat# C6628) or water to inhibit lysosomal degradation for 4 hours.

### RNF13 and RNF167 knockdown

The TriFECTa Dicer Substrate duplex RNAi kit against human RNF13 (#1: 5’rGrGrUrGrArArUrCrArUrCrArGrCrUrArA-3’ and 5’-rUrCrArGrArGrArGrArArUrUrArGrCrUrGrArU-3’, and #3: 5’-rCrGrArCrArUrUrGrArGrGrUrArC-3’ and 5’-rGrUrArGrUrArCrCrUrCrArArUrG-3’) and human RNF167 (#1: 5′-rArArCrUrUrUrGrArCrCrUrCrArArGrGrUrC-3′ and 5′-rCrArUrUrUrArGrGrArCrCrUrUrGrArGrGrU-3′, #2: 5′-rUrCrGrArCrUrUrArCrCrArArArGrArGrCrA-3′ and 5′-rUrUrUrCrArGrUrUrGrCrUrCrUrUrUrGrGrU-3′) was purchased from IDT. The negative control scramble DsiRNA (cat. #DSNC1) was used. Briefly, 2 µL of LipofectAMINE 2000 transfection reagent diluted in 250 µL Extreme MEM was added to a total of 10 nM DsiRNAs diluted in 250 µL Extreme MEM. The mixture was incubated for 20 mins at room temperature and added to the cells. The cells were incubated for 48h and were splitted to be transfected with the appropriate plasmids as described above. Cells were incubated for another 24 hours, for a total of 72 hours of DsiRNA transfection.

### Cell Lysis and Co-immunoprecipitation Assays

Cell lysis was performed as previously described [19]. For co-immunoprecipitation to detect the different form using anti-IDS, 600 µL of total lysate was agitated at 15 rpm for 2h at 4°C with 1 µg of purified anti-HA.11 rabbit polyclonal (BioLegend, Cat# 902302) and 10 µL of Protein A/G Plus-Agarose resin (Santa Cruz Biotechnology, Cat# sc-2003) pre-equilibrated in lysis buffer. Washes were performed with lysis buffer before protein were eluted in 2X Laemmli buffer, boiled for 5 mins at 95°C and loaded on SDS-PAGE. For co-immunoprecipitation to detect the V5, 600 µL of total lysate was agitated at 15 rpm for 2h at 4°C with 10 µL of anti-HA magnetic beads (Bimake.com, Cat# B26201) pre-equilibrated in lysis buffer. After washes with the lysis buffer, proteins were eluted with 1X Laemmli buffer and boiled for 5 mins at 95°C before loading on SDS-PAGE. For co-immunoprecipitation with RNF167/RNF13/IDS, 400 uL of total lysate was agitated at 15 rpm for 2h at 4°C with either 10 uL of anti-HA magnetic beads or with 10 µL of GFP-Catcher beads (Antibodies-online Inc., Limerick, PA, USA, Cat# ABIN5311508) pre-equilibrated in lysis buffer. After washes with the lysis buffer, proteins were eluted with either 1X (anti-HA beads) or 2X (GFP-Catcher beads) Laemmli buffer and boiled for 5 mins at 95°C before loading on SDS-PAGE.

### Glycosylation Assays

45 µL of whole cell lysates were incubated at 100°C for 10 minutes in 1X Glycoprotein Denaturing Buffer (New England Biolabs, Cat# P0703). Subsequently, the proteins were digested with either 1,000 U of EndoHf in 1X GlycoBuffer 3 (New England Biolabs, Cat# P0703) or 500 U of PNGase F in 1X GlycoBuffer 2, which was supplemented with 1% NP-40 (New England Biolabs, Cat# P0704). The digestions were performed overnight at 37°C, after which the reaction were stopped by adding Laemmli buffer. These samples were separated into three distinct SDS-PAGE gels. For the treatment of co-immunoprecipitated protein, the immunoprecipitation was performed as described above. Immunopurified proteins were incubated at 100°C for 10 minutes in 1X Glycoprotein Denaturing Buffer and digested with 500 U of EndoHf in 1X GlycoBuffer 3 overnight at 37°C, before being stopped with Laemmli buffer. 2/3 of the volume of the reactions were loaded on SDS-PAGE for separation to detect using the anti-V5 while 1/3 was detected with the anti-HA.

### SDS-PAGE and Western Blot

For the glycosylation assay only, protein samples were separated on a 1.0 mm Criterion TGX Precast Gel Any kD (BioRad, Cat# 5671124). Otherwise, protein samples were separated on a 1.5 mm SDS-PAGE gel containing 0.5% of 2,2,2-trichloroethanol (TCE). Proteins were transferred to a 0.45 µm PVDF membrane using the Bio-Rad Trans-Blot Turbo system (7 mins for 1.0 mm; 10 mins for 1.5 mm, constant 2.5 A, 25V). Proteins on both gels and membranes were then visualized using the *stain-free* mode of the ChemiDoc MP imaging System (Bio-Rad) and Bio-Rad Image Lab software (except for the precast gels that did not contain TCE). Non-specific sites were blocked using 5% skim milk in TBS-T (20 mM Tris-HCl pH 7.5, 140 mM NaCl, 0.3 % Tween-20) for 1h at room temperature. Membranes were incubated overnight at 4°C with primary antibodies diluted in TBS-T supplemented with 0.05 % NaN_3_. The membranes were washed three times with TBS-T before being incubated for 1h at room temperature with HRP-conjugated secondary antibodies diluted in TBS-T. Immune complexes were revealed using the Clarity Western ECL substrate (BioRad, Cat# 1705060). The ChemiDoc MP imaging system was used to acquire the chemiluminescent signal using automatic time exposure to detect intense bands (short exposure) or automatic time exposure to detect faint bands and/or manual time exposure mode (long exposure).

### Protein Abundance and Statistical Analysis

The relative abundance of different IDS forms from immunoblots was analyzed using short exposure. As IDS immunoblots revealed some mature glycosylated forms and underglycosylated precursor forms at similar MW, the quantifications were performed on the anti-V5 immunoblots. The mature and precursor forms were detected using the anti-IDS, which detected only the N-terminal SD1. The band signal obtained by immunoblotting against V5 or IDS was normalized against the corresponding stain-free lane, and the results were then normalized against the appropriate control lane. Three independent, non-blinded experiments were performed for all assays. Datasets were exported as a CSV file and collected in Microsoft Excel before being subjected to statistical analysis in GraphPad Prism 10. Parametric tests were used due to the Gaussian distribution of the data. When two groups were compared, statistical difference was determined using a two-tailed paired t-test. When multiple groups with one grouping variable were compared, statistical difference was determined by a repeated measures one-way ANOVA with Tukey’s multiple comparisons test. When multiple groups with two grouping variables were compared, statistical difference was determined using a repeated measures two-way ANOVA with uncorrected Fisher’s LSD or Dunnett’s multiple comparisons test to compare means with others in its row and its column.

### Immunostaining

Transfected HEK293T/17 or HeLa cells were washed three times in PBS before being fixed for 15 mins with 4% paraformaldehyde (Electron Microscopy Sciences, Cat# 15710) and 4% sucrose in PBS at 37°C (HEK293T/17) or room temperature (HeLa). Fixed cells were washed three times and permeabilized with 0.25% Triton X-100 in PBS for 15 mins at room temperature. Non-specific sites were blocked using a blocking solution composed of 10% normal goat serum (NGS) (+ 0.1% Triton X-100 for HEK293T/17 cells) in PBS for 1 hour. Subsequent staining was performed by incubating with primary antibodies diluted in 5% NGS (+ 0.05% Triton X-100 for HEK293T/17 cells) in PBS overnight at 4°C (HEK293T/17) or 1h at room temperature (HeLa). Coverslips were washed three times with PBS, followed by incubation with the appropriate conjugated secondary antibodies diluted in 5% NGS (+ 0.05% Triton X-100 for HEK293T/17 cells) in PBS for 1 hour at room temperature. After being washed in PBS, the coverslips were mounted using ProLong Diamond Antifade (Thermo Fisher Scientific Cat# P36961) and allowed to dry for 2 hours at 37°C. The mounted coverslips were then stored at 4°C until image acquisitions were performed.

### Fluorescence Microscopy Image Acquisition

Image acquisition was carried out using an inverted Olympus IX83 epi-fluorescence microscope equipped with a U plan S-Apo 60x / 1.35 numerical aperture oil objective (Olympus), an X-Cite Xylis 365 LED-based illumination source (Excelitas Technologies Corp.), and a Zyla 4.2 Plus sCMOS camera (Andor). The filters and dichroic mirror employed by this widefield system have been thoroughly described in [19]. Image acquisition was facilitated by using the Olympus CellSens Dimension software version 2.2 (Olympus, RRID: SCR_014551). The resolution was set at 2048 x 2048 pixels, and z-stack images were captured at 0.27 µm intervals for 8 to 10 optical slices. Lamp intensity and exposure time were configured to maximize signal intensity without reaching saturation and maintained constant within an independent experiment. Prior to analysis, deconvolution was performed on the images using the Olympus 3D Deconvolution feature in the Olympus CellSens Dimension software.

### Image Analysis and Statistical Analysis

Colocalization assays were analyzed as previously described [19]. Three independent, non-blinded experiments were performed for all assays. The resulting image analysis datasets were exported to a CSV file and collected in Microsoft Excel before being undergoing statistical analysis in GraphPad Prism 10. Due to the Gaussian distribution of the data, parametric tests were used. When two groups were compared, statistical significance was determined using a two-tailed paired t-test. When multiple groups with one grouping variable were compared, statistical significance was determined by a repeated measures one-way ANOVA with Tukey’s multiple comparisons test. When multiple groups with two grouping variables were compared, statistical difference was determined using repeated measures two-way ANOVA with uncorrected Fisher’s LSD multiple comparisons test to compare means with others in its row and its column.

### Predicted Structure

The structure of the complexes formed by RNF13 (Uniprot accession O43567), RNF137 (Uniprot accession Q9H6Y7) and IDS (Uniprot accession P22304) and its glycosylated form were modeled using the AlphaFold 3 server [52]. The RNF13–IDS complex was visualized using the PyMOL Molecular Graphics System, Version 3.1 Schrödinger, LLC.

## ACKNOWLEDGEMENTS

This work is part of the requirements for VCC’s Ph.D. thesis at UQAM. The CERMO-FC funded this study via its graduate scholarship *Félix-Antoine Aublet Research Grant* awarded to VCC and AMS in 2023. PhD graduate scholarship from FRQNT were awarded to VCC (DOI: doi.org/10.69777/326578) and AYB (DOI: doi.org/10.69777/317082). Funding sources had no role in study design, data collection, analysis, or interpretation.

## ABBREVIATIONS

AP: Adaptor protein;
CHX: Cycloheximide;
CQ: Chloroquine;
DN: Dominant negative;
EGAD: Endosome and Golgi-associated degradation;
Endo H: Endoglycosidase H;
ER: Endoplasmic reticulum;
ERAD,: ER-associated degradation;
GAG: Glycosaminoglycan;
IDS: Iduronate 2-sulfatase;
M6P: Mannonse-6-phosphate;
M6PR: M6P receptor;
MOC: Manders’ overlap coefficient
MPS II: Mucopolysaccharidosis type II;
MW: Molecular weight;
PA: Protease-associated;
PNGase F: Peptide:N-glycosidase F;
RING: Really Interesting New Gene;
Tg: Thapsigargin;
TGN: Trans-Golgi Network;
TM: Transmembrane;
Tn: Tunicamycin

## CONFLICT OF INTEREST

No competing interests are declared.

## Notes

### Competing Interest Statement

The authors have declared no competing interest.

